# Environment-Dependent Switch between Two Immune Systems against *Ralstonia solanacearum* Infections in Pepper

**DOI:** 10.1101/2021.01.14.426660

**Authors:** Sheng Yang, Weiwei Cai, Lei Shen, Ruijie Wu, Jianshen Cao, Jinsen Cai, Shicong He, Yutong Zheng, Qixiong Zhang, Aiwen Wang, Deyi Guan, Shuilin He

## Abstract

- Although plant diseases generally cause more severe symptoms under conditions of high temperature and high humidity(HTHH), how plant respond to pathogen attack under this condition remains elusive.
- As an example, we herein comparatively studied pepper(*Capsicum annuum*) immunity against *Ralstonia solanacearum* under HTHH and ambient temperature by approaches of reverse genetics. We found that pepper respond to *R*.*solanacearum* infection by activating salicylic-acid- and jasmonic-acid-mediated immunity at ambient temperature. Under HTHH, However, it no longer activates JA-mediated immunity and activates only transient SA signaling at the early stage of *R*.*solanacearum* infection, but instead induces cytokinin mediated immunity.
- These two immune systems are positively regulated by CaWRKY40 via binding the WT-box and a novel W-box like (WL) box, respectively, in an environment-dependent manner: *CaWRKY40* is activated upon *R. solanacearum* infection under HTHH, thereby upregulating ISOPENTENYLTRANSFERASE5(IPT5). The resulting cytokinin then works synergistically with CaWRKY40 in activating a subset of glutathione S-transferase genes via chromatin activation and WL-box binding, but prevent CaWRKY40 from activating ISOCHORISMATE SYNTHASE1 (ICS1) or SA-/JA-dependent pathogenesis-related genes by chromatin inactivation or by blocking WT-box binding.
- These results highlight the specific pepper immune response to *R. solanacearum* infection under HTHH and its synergistic activation by *Ca*WRKY40 and cytokinins.

## Introduction

Recurring stresses, alone or in combination, can exert constraints on plant evolution and shape how plant traits evolve. Plants have evolved a sophisticated immune system comprising two interconnected branches: effector-triggered immunity (ETI) and pathogen-associated molecular pattern (PAMP)-triggered immunity (PTI). During PTI and ETI, pathogen-derived PAMPs and effectors are perceived by plant-membrane-localized pattern-recognition receptors (PRRs) and intracellular resistance (R) proteins, respectively (Jones & Dangl, 2006). Upon pathogen attack and activation of these two layers of immunity, plants produce the anti-microbial compounds phytoalexins (Chang *et al*., 2017) and pathogenesis-related (PR) proteins (Liu *et al*., 2017; Silva *et al*., 2018) with different timing and magnitude. To respond to pathogens appropriately, PTI and ETI are tightly regulated by partially overlapping signaling pathways mediated by phytohormones including salicylic acid (SA), jasmonic acid (JA), ethylene (ET), abscisic acid (ABA), and cytokinins (CKs) (Robert-Seilaniantz *et al*., 2011). Immune signaling components can be specific for pathogens of different lifestyles; for instance, SA-mediated signaling is generally involved in immunity to biotrophic pathogens, while JA participates in responses to necrotrophic pathogens (Glazebrook, 2005). CKs also promote immunity, although biotrophic pathogens benefit from a CK-mediated increase of sink organ strength, which enhances nutrient supply to infected leaves (Walters & McRoberts, 2006). CKs, including the natural biologically active compound *trans*-zeatin (tZ), contribute to immunity against hemi-biotrophic or biotrophic pathogens (Grosskinsky *et al*., 2013; Reusche *et al*., 2013; Albrecht & Argueso, 2017). However, the roles of CKs in plant immunity seem complicated: in some cases they act negatively in response to pathogens (Wasilewska *et al*., 2008; Meng *et al*., 2013; Naseem *et al*., 2014; Su *et al*., 2017). The causes of these inconsistent activities are unclear.

Transcription factors (TFs) integrate signals from the plant immune system for transcriptional reprogramming by targeting specific sets of responsive promoters (Tsuda & Somssich, 2015). The WRKY TF family in plants is implicated in biological processes ranging from growth and development to defense against abiotic and biotic stresses, (Pandey & Somssich, 2009). Upon pathogen attack, many WRKY genes are upregulated and form WRKY-dependent networks, leading to immune response outputs (Eulgem & Somssich, 2007). WRKY TFs bind specifically to their cognate *cis* elements within target promoters. Most functionally characterized WRKY TFs bind a typical W-box motif (TTGACC/T), although a few also bind a WT-box motif (ATGCCC; (Machens *et al*., 2014; Kanofsky *et al*., 2017). The specific binding of WRKY TFs to their target genes via W-box is partially affected by the adjacent bases (Ciolkowski *et al*., 2008) and by interacting proteins (Chi *et al*., 2013). However, the roles of other *cis* elements in the functional specificity of WRKY TFs remains under-investigated.

Plant–pathogen interactions are generally affected by the environment. Plants frequently experience high temperature and high humidity (abbreviated HTHH here) together, which influences the outcome of plant–pathogen interactions, particularly in the Solanaceae family during infection by soil-borne pathogens. High temperatures (Hwang *et al*., 2000; Xiao *et al*., 2003; Bieri *et al*., 2004) or high humidity (Jambunathan *et al*., 2001; Zhou *et al*., 2004; Noutoshi *et al*., 2005) attenuate R protein– mediated plant immunity, exacerbating plant disease symptoms. The mechanisms of plant immunity and thermotolerance have been intensively investigated, but most studies have focused on a single stress. As plant responses to multiple concomitant stresses may differ from those to individual stresses (Atkinson & Urwin, 2012), how plants respond to pathogen attacks under HTHH remains poorly understood. Pepper is a solanaceous vegetable of agricultural importance worldwide that is devastated by bacterial wilt, a soil-borne disease caused by the bacterium *Ralstonia solanacearum*s (Mansfield *et al*., 2012). Bacterial wilt symptoms are generally more severe under HTHH than under ambient temperatures, such that only pepper cultivars with high resistance to bacterial wilt under HTHH will survive an attack by R. *solanacearum*. The pepper TFs CaWRKY6 and CaWRKY40 promote pepper immune responses to HTHH and to *R. solanacearum* (Dang *et al*., 2013; Cai *et al*., 2015), suggesting a close relationship between pepper responses to heat stress and pathogen infection.

Here, we demonstrate that pepper employs SA- and JA-mediated immunity to cope with *R. solanacearum* infections at ambient temperatures, but switches to CK-mediated defenses to protect itself against *R. solanacearum* infection under HTHH (abbreviated RSHT). Our data establish that Ca*WRKY40* is activated upon RSHT, thereby inducing expression of the CK-biosynthetic gene *ISOPENTENYLTRANSFERASE 5* (*IPT5*). The resulting increase in CK in plant tissues confers immunity against RSHT through upregulation of *PR* genes including *MICROSOMAL GLUTATHIONE S-TRANSFERASE 3* (*Mgst3*) and *PROLINE-RICH PROTEIN 1* (*PRP1*), via synergistic action of CK signaling and CaWRKY40, which prevents the activation of SA signaling by CaWRKY40.

## Materials and Methods

### Plant Materials and Growth Conditions

We used the pepper (*Capsicum annuum*) inbred lines HN42 and TT5203, as well as tobacco (*Nicotiana tabacum* cv. Cuibi 1), *Nicotiana benthamiana*, and tomato (*Solanum lycopersicum* cv. Micro-Tom) plants, in the present study. We also generated *CaWRKY40*-overexpressing lines in the pepper HN42 and TT5203 inbred lines and in *N. benthamiana*; *CaIPT5*-overexpressing *N. benthamiana* lines; and silencing lines against *CaWRKY40* and *CaIPT5* in the HN42 pepper and *CaIPT5*-overexpressing *N. benthamiana* backgrounds.

Plants were grown on peat soil in plastic pots kept in growth chambers under the conditions of 27–29°C, 60–70 µmol photons m^-2^ s^-1^, 60% relative humidity, and a 16 h light/8 h dark cycle. When investigating the effect of exogenous application of the phytohormones SA, methyl jasmonate (MeJA), tZ, KT, or 6-BA on pepper responses to *R. solanacearum* infection, we grew seedlings on Murashige and Skoog (MS) medium and placed them in the growth room under the conditions described above.

### RSRT, RSHT, HTHH, and RTHH Treatments

Soil-grown pepper, tobacco, and tomato plants had their roots mechanically damaged before inoculation via *R. solanacearum* by root irrigation with 5 mL (per plant) of a cell suspension of *R. solanacearum* strain GMI1000 (10^8^ cfu/mL). The plants were placed in an illuminated incubator (60–70 µmol photons m^-2^ s^-1^,16-h light/8-h dark photoperiod) at either 28°C (for *R. solanacearum* infection at room temperature, RSRT treatment) or 37°C (for *R. solanacearum* infection at high temperature, RSHT treatment). The soil in the pots was kept at its maximum water holding capacity, while air humidity was kept at least at 80%. Non-inoculated plants were grown under the same conditions with mechanical root damage but without bacterial inoculation, either at 28°C (room temperature, high humidity; RTHH treatment), or 37°C (high temperature, high humidity; HTHH treatment).

For ChIP and ChIP-seq assays, we infiltrated the leaves of pepper plants to transiently overexpress *CaWRKY40*, or the roots of stable *CaWRKY40* transgenic pepper plants with *R. solanacearum* (OD_600_ = 0.4) with a disposable sterilized syringe. The *R. solanacearum* suspension was infiltrated near the main leaf vein and expanded through the leaf to the other side of the main vein.

### ChIP

ChIP assays were performed according to the published protocol by Khan *et al*. (2018). We collected leaves or roots of pepper plants at the indicated times and crosslinked the tissues in 1% (w/v) formaldehyde 5min on ice, after which we isolated chromatin and sheared the genomic DNA into fragments of 300–500 bp in length. We then immunoprecipitated the DNA-protein complexes using anti-GFP, anti-H3K4me3, or anti-H3K9me2 antibodies. We then reversed the crosslinking and purified the DNA released from DNA-protein complexes. The DNA was used as template for PCR or quantitative PCR (qPCR) using specific primer pairs (Table S1). The ChIP-seq data sets used in this study have been deposited at the CNSA repository(CNP0001155).

### ChIP-seq Analysis

We infiltrated 30 fully expanded leaves from pepper plants that had been challenged by RTHH, RSRT, HTHH, or RSHT treatment at the 6-leaf stage with Agrobacterium cell suspensions harboring the binary vector *p35Spro:CaWRKY40-GFP*. After 48 h, the infiltrated leaves were harvested and mixed at 1:1:1:1 ratio and were subjected to ChIP following the above-mentioned method. We subjected the purified genomic DNA fragments to linear DNA amplification (LinDA) in order to generate sufficient material to construct ChIP-seq sequencing libraries using a NEBNext® ChIP-seq Library Pre Reagent Set for Illumina® (New England Biolabs, Ipswich, MA). DNA sequencing was performed on an Illumina Hiseq2500 platform (Novogene, Beijing, China) and resulted in about 10 million 100 bp single-end reads per sample. We removed low quality reads, those with over 15% ambiguous bases, reads contaminated with 5’ barcode, and reads without 3’ linker sequences or inserts, trimmed the 3’ linker sequences, and discarded reads shorter than 18 nt after data cleaning. The remaining reads were aligned to the pepper reference genome by BWA (Burrows Wheeler Aligner) (H & R, 2009). We used the MACS2 software to predict DNA fragment sizes, which we then used for subsequent peak analysis. We also used the MACS2 software (with threshold *q*-value = 0.05) to detect signal peaks, as well as for analysis of the number, width, and distribution of peaks, and the underlying genes identified by the peaks (Y *et al*., 2008). The TSS (transcription start site) of each peak-related gene was detected by Peak Annotator (M *et al*., 2010). To search for conserved binding motifs across CaWRKY40 binding regions, we extracted 500 bp of genomic sequence surrounding all peak maxima and submitted the sequences to the online version of MEME-ChIP (P & TL, 2011).

### RT-qPCR

We performed RT-qPCR assays as previously described (Dang *et al*., 2013). Briefly, total RNA was extracted from the roots or leaves of pepper or *N. benthamiana* plants using Trizol reagent (Invitrogen, Carlsbad, CA, USA). Reverse transcription was carried out using the HiScript III RT SuperMix (R323-01, Vazyme Biotech, China). qPCR was carried out by using SYBR Green (ThermoFisher Scientific, USA) according to the manufacturer’s instructions. Four independent biological duplicates were generally performed with appropriate specific primers (Table S1). Expression values were normalized to *CaACTIN* expression.

### Supporting information for materials and methods

Details of the experimental programs for Vector Construction (Methods S1), Genetic Transformation of Pepper and *N. benthamiana* (Methods S2), Virus-Induced Gene Silencing (Methods S3), Exogenous Applications of SA, JA, tZ, KT, or 6-BA (Methods S4), RNA-seq Analysis(Methods S5), EMSA Analysis(Methods S6), MST Analysis(Methods S7), GUS Activity Assay(Methods S8), prokaryotic Expression and Protein Purification(Methods S9), plant Total Protein Extraction and Immunoblot Analysis(Methods S10), phytohormone Measurements and Quantification(Methods S11).

## Results

### RSRT and RSHT Produce Varying Effects in Different Pepper Inbred Lines

To test if pepper resistance to bacterial wilt was compromised by HTHH, we inoculated 25 pepper inbred lines with *R. solanacearum* via root irrigation and exposed them to room temperature high humidity conditions (RTHH; 28°C, air humidity >80%, with inoculation denoted RSRT) and HTHH (37°C, air humidity >80%, with inoculation denoted RSHT). As controls, the same pepper inbred lines were exposed to RTHH or HTHH but not inoculated with *R. solanacearum*. HTHH alone caused no obvious damage to the tested pepper lines. By contrast, most inoculated lines exhibited more severe bacterial wilt symptoms and supported higher *R. solanacearum* growth, based on the number of colony-forming units (cfu), under HTHH compared to RTHH (Fig. S1). Notably, some pepper inbred lines displayed no obvious bacterial wilt symptoms upon RSHT even at 10 d post inoculation (dpi). In those lines, bacterial wilt resistance was attenuated by HTHH, which may occur broadly in the Solanaceae, since resistance of *Nicotiana benthamiana* and tomato (*Solanum lycopersicum*) to *R. solanacearum* inoculation was similarly impaired by HTHH (Fig. S2).

### RSHT and RSRT Activate Different PR Genes

To determine the genes differentially expressed in pepper plants under RSHT and RSRT, we profiled the transcriptome of plant roots from the TT5203 (more susceptible to RSHT) and HN42 (less susceptible to RSHT) pepper inbred lines challenged with RSRT and RSHT. We focused on the root transcriptome, as *R. solanacearum* invades pepper plants exclusively through this tissue (Digonnet *et al*., 2012). We mechanically damaged the roots of seedlings before dipping them into a solution of *R. solanacearum* cells (OD_600_ = 0.6) and placing them into HTHH or RTHH conditions, using non-inoculated seedlings as controls. We harvested samples for RNA isolation and deep sequencing of the transcriptome (RNA-seq analysis) at nearly the same stage of pathogenesis to minimize variation in the extent of infection. We collected the roots of TT5203 pepper plants under RSHT and RSRT at 90 h post inoculation (hpi) and 48 hpi, respectively, while we harvested HN42 roots at 90 hpi and 24 hpi, respectively. The harvested plants all showed similar bacterial wilt symptoms (Fig. 1a). Non-inoculated TT5203 and HN42 pepper plants under RTHH or HTHH were harvested at the same time points as the corresponding inoculated plants exposed to HTHH.

**Fig. 1.**
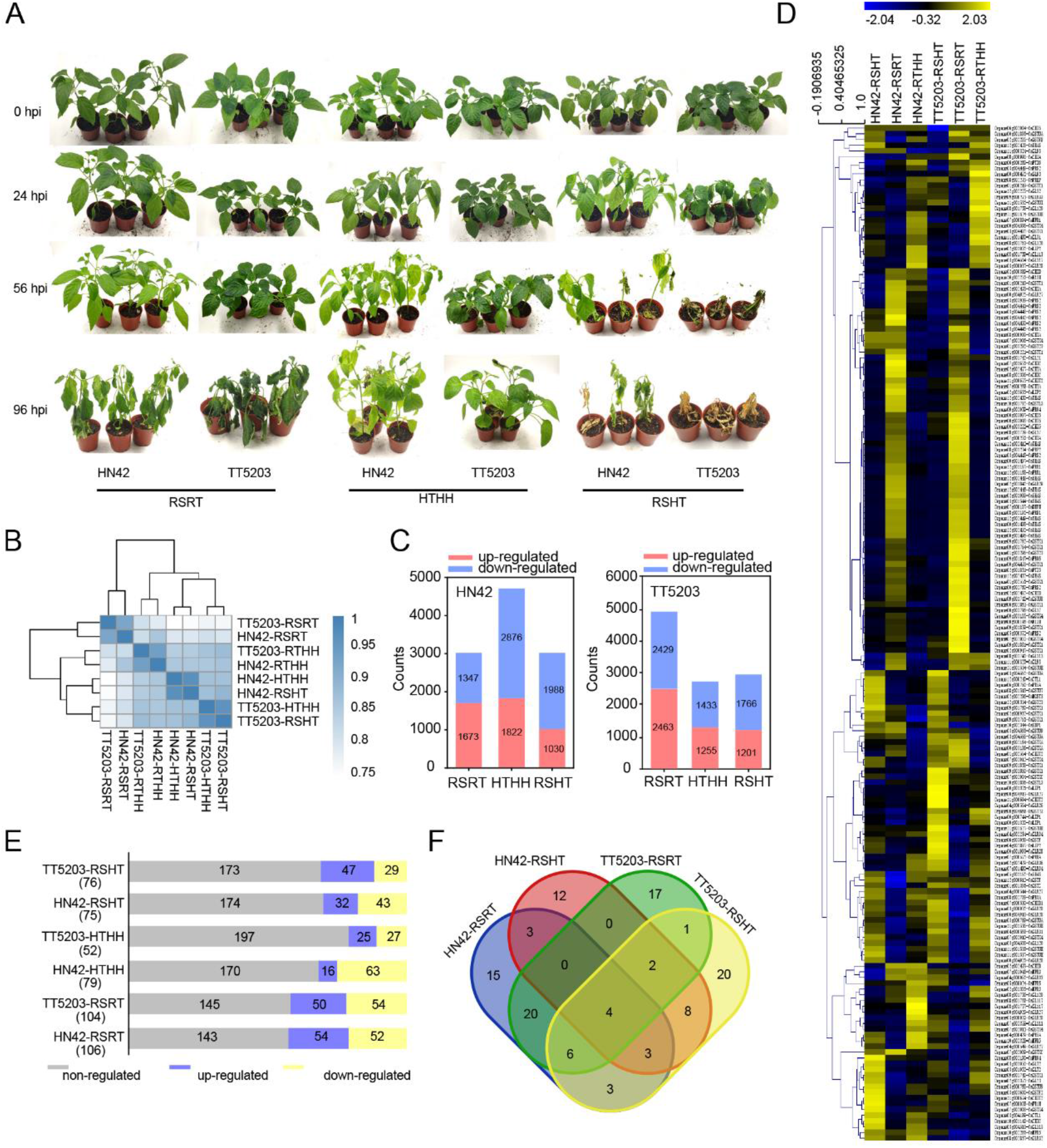
Comparative Transcriptome Analysis of Different Pepper Inbred Lines in Response to RSHT and RSRT. (a) Phenotypes of TT5203 and HN42 pepper inbred lines upon RSRT, HTHH, or RSHT. (b) Correlations among the transcriptomes of the TT5203 and HN42 pepper inbred lines upon RTHH, RSRT, HTHH, or RSHT by R2 matrix heatmap and cluster detection. (c) Genes induced in the TT5203 and HN42 pepper inbred lines by RSRT, HTHH, or RSHT. Red and blue bars indicate the total numbers of genes up- or down-regulated, respectively. (d) Transcript levels of *PR* genes upon RSRT, HTHH, or RSHT in the TT5203 and HN42 pepper inbred lines, shown as a heatmap. In the color scale, blue, black, and yellow indicate low, intermediate, and high expression levels, respectively. (e) Numbers of *PR* genes up- or down-regulated by RSRT, HTHH, or RSHT in the roots of the TT5203 and HN42 pepper inbred lines. The numbers of differentially expressed genes are shown in parentheses. (f) Overlap of *PR* genes upregulated by RSRT and RSHT in the TT5203 and HN42 pepper inbred lines. In (b) to (d), three independent biological replicates were analyzed for RNA-seq (*P* < 0.1).

Based on cluster analysis, the transcriptome of RSRT plants strongly correlated with that of RTHH plants, while the transcriptome of RSHT plants correlated with that of HTHH plants (Fig. 1b). Compared to control plants, we identified 3,020 (for RSHT) and 2,968 (for RSRT) differentially expressed genes in the HN42 inbred line, and 4,982 (for RSHT) and 2,967 (for RSRT) differentially expressed genes in TT5203 plants. By contrast, HTHH resulted in 4,698 (in HN42) and 2,688 (in TT5203) differentially expressed genes, with a fold change >2 and *P* value <0.05 (Fig. 1c). Functional annotations of upregulated genes included 75 and 76 defense-related genes (DRGs) after RSHT treatment, and 104 and 106 after RSRT treatment, in the HN42 and TT5203 inbred lines, respectively (Fig. 1d). In addition, 15 and 12 DRGs specifically responded to RSRT and RSHT in HN24, while 17 and 20 DRGs responded specifically to RSRT and RSHT in TT5203, respectively (Fig. 1e). Although genes encoding R proteins were differentially expressed upon RSHT, RSRT, HTHH, and RTHHs in both HN42 and TT5203 inbred lines (Fig. S3), their exact functions in pepper remain unidentified. We thus focused our analysis on genes encoding functionally conserved and well-characterized pathogenesis-related (PR) genes.

Genes encoding endochitinases (Kuc, 1990), the SA-dependent proteins PR1, PR2, and PR5 (Thomma *et al*., 1998), a JA-dependent defensin-like protein (O’Donnell *et al*., 2003), and glutathione S-transferases (GSTs) (Andaya & Tai, 2006; Gong *et al*., 2018) were differentially modulated by RSRT or RSHT in the two inbred lines and were classified into three groups. Group I comprised genes upregulated specifically upon RSRT; group II genes were upregulated upon RSHT in the two tested inbred lines but with higher levels in HN42; and group III genes were upregulated specifically in HN42 plants under RSHT (Fig. 1f). Since HN42 was less susceptible to RSHT, we hypothesized that group II and III genes might contribute to pepper resistance to RSHT.

### RSHT and RSRT Lead to Accumulation of Hormones and Hormone Biosynthetic Gene Transcripts

We examined phytohormone biosynthetic gene transcripts in our RNA-seq datasets to investigate their possible involvement in pepper responses to RSHT. The JA biosynthetic gene *LINOLEATE 9S-LIPOXYGENASE* (*LOX1*) (Spoel *et al*., 2003) and the SA biosynthetic gene *ISOCHORISMATE SYNTHASE1* (*ICS1*) (Attaran *et al*., 2009) were highly upregulated by RSRT, but downregulated by RSHT, in the two tested inbred lines. By contrast, genes encoding enzymes for ABA and CK biosynthesis, such as 9-*cis*-EPOXYCAROTENOID DIOXYGENASE 1 (NCED1), NCED5 (Cutler & Krochko, 1999), ISOPENTENYL TRANSFERASE 2 (IPT2) (Sakakibara, 2005), IPT5, and IPT9, were downregulated by RSRT, but highly upregulated by RSHT (Fig. 2a–2b). These data suggest that SA, JA, ABA, and CK are differentially modulated by RSHT and RSRT.

**Fig. 2.**
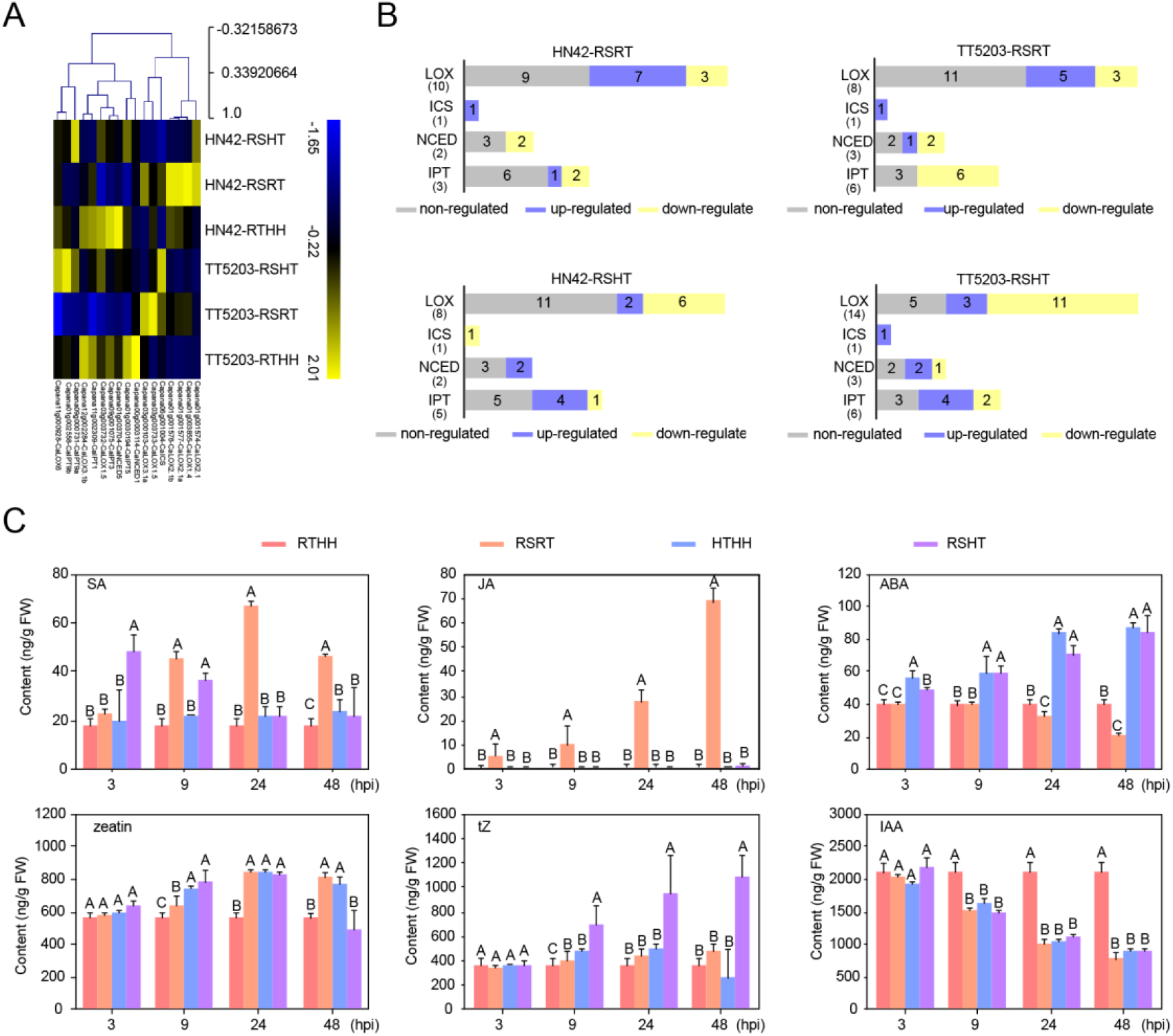
Transcript Levels of Phytohormone Biosynthetic Genes and Phytohormone Levels in the Roots of Pepper Plants Challenged by RSHT or RSRT. (a) Transcript levels of hormone biosynthetic genes upon RTHH, RSRT, HTHH, and RSHT in the roots of TT5203 and HN42 pepper inbred lines, visualized as a heatmap. In the color scale, blue, black, and yellow indicate low, intermediate, and high levels, respectively. (b) Numbers of up- or down-regulated hormone biosynthetic genes upon RTHH, RSRT, HTHH, or RSHT in the roots of TT5203 and HN42 pepper plants. (c) Phytohormone contents for SA, JA, ABA, ZT, tZ, and IAA, measured by LC-MS/MS at different time points in the roots of TT5203 and HN42 pepper plants challenged by RTHH, RSRT, HTHH, or RSHT. Data represent the mean ± SD of three biological replicates. Different capital letters above the bars indicate significant differences (*P* < 0.01) by Fisher’s protected LSD test.

Accordingly, we measured endogenous SA, JA, ABA, and CK levels in the roots of HN42 plants challenged by RSRT and RSHT at 3, 9, 24 and 48 hpi. At 3 and 9 h post treatment (hpt). The roots of RSHT-challenged plants had significantly lower contents of all phytohormones than those of the RSRT plants with the exception of SA, which accumulated to higher levels at these early time points. ABA and tZ, one of the most active CKs, were significant higher in RSHT roots of both inbred lines relative to control plants (Fig. 2c), indicating that RSHT suppressed JA and SA biosynthesis, while promoting CK and ABA biosynthesis.

### *WRKY* Family Members Are Differentially Expressed upon RSHT or RSRT

To identify WRKY TFs involved specifically in pepper responses to RSHT, we searched for WRKY genes that were upregulated in the more resistant HN42 inbred line upon RSHT. In all, RSHT and RSRT induced 38 *WRKY* genes in HN42 or TT5203 (Fig. S4a). Of those, 9 and 13 were upregulated in HN42 and TT5203 plants under RSHT, respectively (Fig. S4b). Only three (*CaWRKY15, CaWRKY22*, and *CaWRKY40*) had higher levels in HN42 than in TT5203, with *CaWRKY40* exhibiting the highest levels (Fig. S4c). For validation, we measured *CaWRKY40* transcript by real-time quantitative PCR (RT-qPCR) in HN42 roots under RSRT or RSHT from 1 to 48 hpi. *CaWRKY40* was upregulated by both treatments (Fig. S4d).

### CaWRKY40 Is a Positive Regulator of Responses to RSHT

Since CaWRKY40 promotes both resistance to RSRT and thermotolerance in pepper (Dang *et al*., 2013), we hypothesized that CaWRKY40 also promotes resistance to RSHT. Virus-induced gene silencing (VIGS) of *CaWRKY40* increased the susceptibility of pepper plants to RSHT and RSRT, based on the bacterial wilt symptom severit at 48 hpi (Fig. 3a–3c). By contrast, HN42 and TT5203 plants overexpressing *CaWRKY40* had enhanced resistance to RSRT and RSHT (Fig. 3d–3e), with lower disease indices (Fig. 3f) and less *R. solanacearum* growth (Fig. 3g). Heterologous expression of *CaWRKY40* in *N. benthamiana* plants produced similar effects (Fig. 3h– 3k). Thus, CaWRKY40 promotes resistance to RSRT and RSHT.

**Fig. 3.**
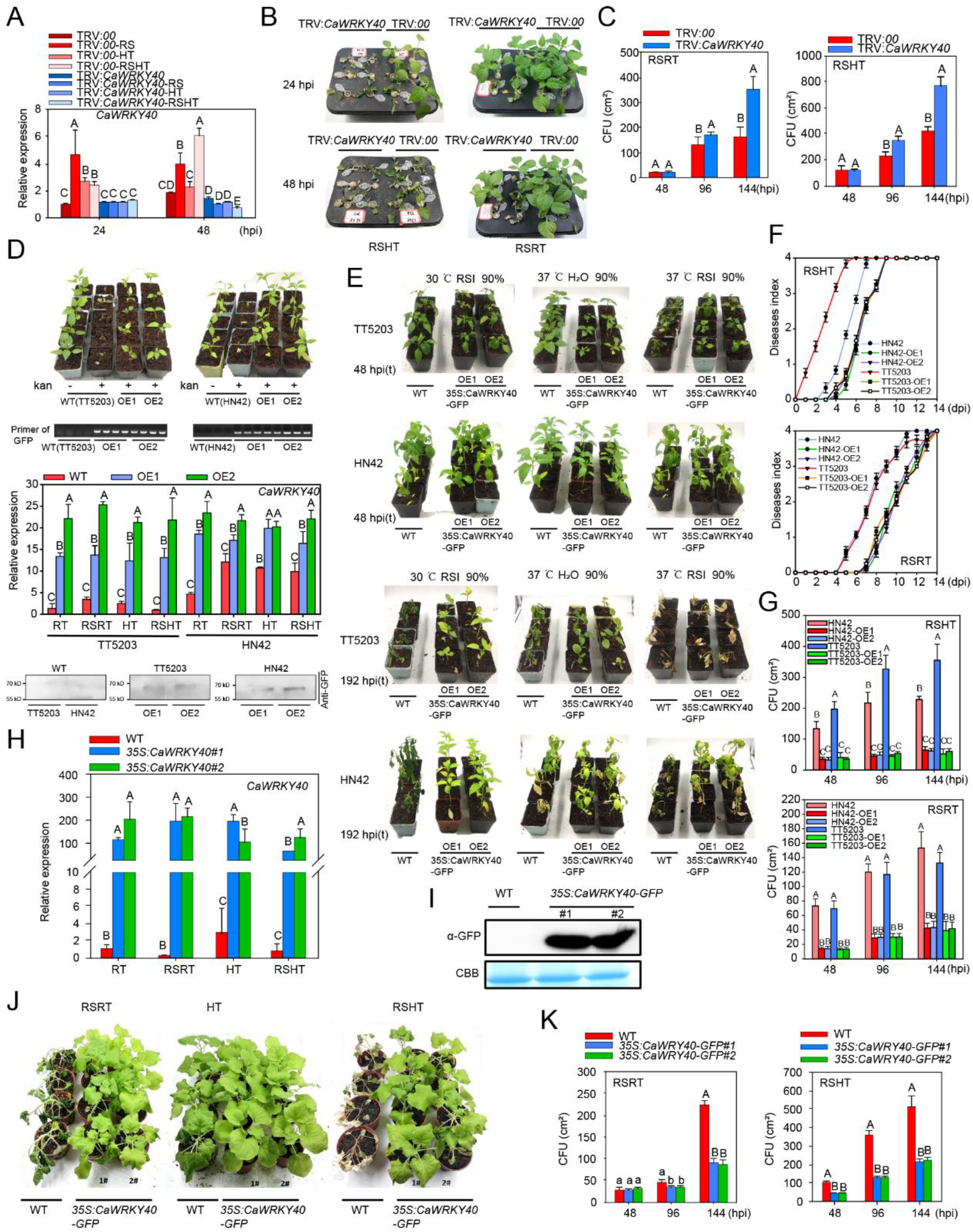
CaWRKY40 Functions Positively in Pepper Response to RSHT. (a) Successful silencing of *CaWRKY40* in pTRV:*CaWRKY40* pepper plants, as shown by RT-qPCR in the roots of pTRV:*CaWRKY40* plants challenged with RTHH, RSRT, HTHH, or RSHT treatment at 24 and 48 hpt. (b) Bacterial wilt symptoms of pTRV:*CaWRKY40* HN42 pepper plants and their control plants to RSRT and RSHT treatment at 3dpt. (c) Growth of *R. solanacearum*, expressed as *R. solanacearum* cfu, in leaves of pTRV:*CaWRKY40* HN42 plant and their control plants challenged with RSRT and RSHT treatment at 48, 96, and 144 hpi. (d) Selection and confirmation of TT5203 and HN42 pepper plants overexpressing *CaWRKY40* with kanamycin treatment, PCR amplification, and RT-qPCR with specific primer pairs as well as immunoblot analysis using an anti-GFP antibody. The kanamycin solution (50 mg/L) was sprayed onto pepper plant roots to select transgenic pepper plants. Positive TT5203 and HN42 pepper plants were screened by PCR using specific primers listed in supplemental tables. Expression of *CaWRKY40-GFP* in TT5203 and HN42 non-transgenic and *CaWRKY40* transgenic pepper plants was determined by RT-qPCR (transcript) and immunoblot analysis (protein). (e) Bacterial wilt symptoms of TT5203 and HN42 plants overexpressing *CaWRKY40* and challenged with RSRT, HTHH, or RSHT treatment at 48 hpt and 192 hpt. RSI: *R. solanacearum* inoculation. (f) Disease index of TT5203 and HN42 plants overexpressing *CaWRKY40* and challenged with RSRT, HTHH, or RSHT treatment, from 0 to 14 dpi. Disease index score was determined according to the percentage of wilted leaves observed in pepper plants. The disease index values were defined as: 0 (no symptoms), 1 (>0–25% wilted leaves), 2 (25–50%), 3 (50–75%), and 4 (75–100%). (g) *R. solanacearum* growth (cfu) in leaves of TT5203 and HN42 plants overexpressing *CaWRKY40* and their control plants, challenged with RSHT or RSRT treatment at 48, 96, and 144 hpt. (h) and (i) Expression of *CaWRKY40* in leaves of *N. benthamiana* plants overexpressing CaWRKY40-GFP and challenged with RTHH, RSRT, HTHH, or RSHT treatment, as seen by RT-qPCR (for transcript) and immunoblot analysis (protein). (j) Bacterial wilt symptoms of *N. benthamiana* plants overexpressing CaWRKY40-GFP and their control plants challenged with RSRT or RSHT treatment at 3 dpt. (k) *R. solanacearum* growth (cfu) in leaves of *N. benthamiana* plants overexpressing CaWRKY40-GFP and their control plants, challenged with RSRT or RSHT treatment at 48, 96, and 144 hpt. In (a), (c), (d), (g), (h), and (k), data represent the mean ± SD of three biological replicates. Different capital letters above the bars indicate significant difference (*P* < 0.01), determined using Fisher’s protected LSD test.

### CaWRKY40 Functions in Pepper Immunity to RSRT and RSHT via Differential Targeting

To identify the direct target genes of CaWRKY40, we performed Chromatin immunoprecipitation sequencing (ChIP-seq), using chromatin from the roots of pepper plants overexpressing *CaWRKY40-GFP* and challenged with RSHT or RSRT. We identified 5,766 CaWRKY40 binding sites associated with 1,434 genes. We also performed RNA-seq on RNA isolated from *CaWRKY40*-silenced pepper plants under RSHT to identify genes regulated by CaWRKY40 upon RSHT. We identified candidate CaWRKY40 direct target *PR* genes via comparison of the genes upregulated in HN42 plants under RSHT, those downregulated in *CaWRKY40*-silenced HN42 plants under RSHT, and CaWRKY40 targets from ChIP-seq (Table S1; Fig. 4b–4c). CaWRKY40 targeted and regulated the expression of *SALT TOLERANCE HOMOLOG2* (*STH2*) and *DEFENSELESS1* (*DEF1*) under RSRT, while regulating *Mgst3* and *PRP1* upon RSHT (Fig. 4d). In addition, *Mgst3* and *PRP1* transcript levels correlated with RSHT resistance (Fig. 4e), with more resistant lines characterized by higher expression of these two genes. Thus, specific upregulation of *Mgst3* and *PRP1* under RSHT may occur in various pepper lines and a distinct defense responses occur in pepper plants under RSHT and RSRT.

**Fig. 4.**
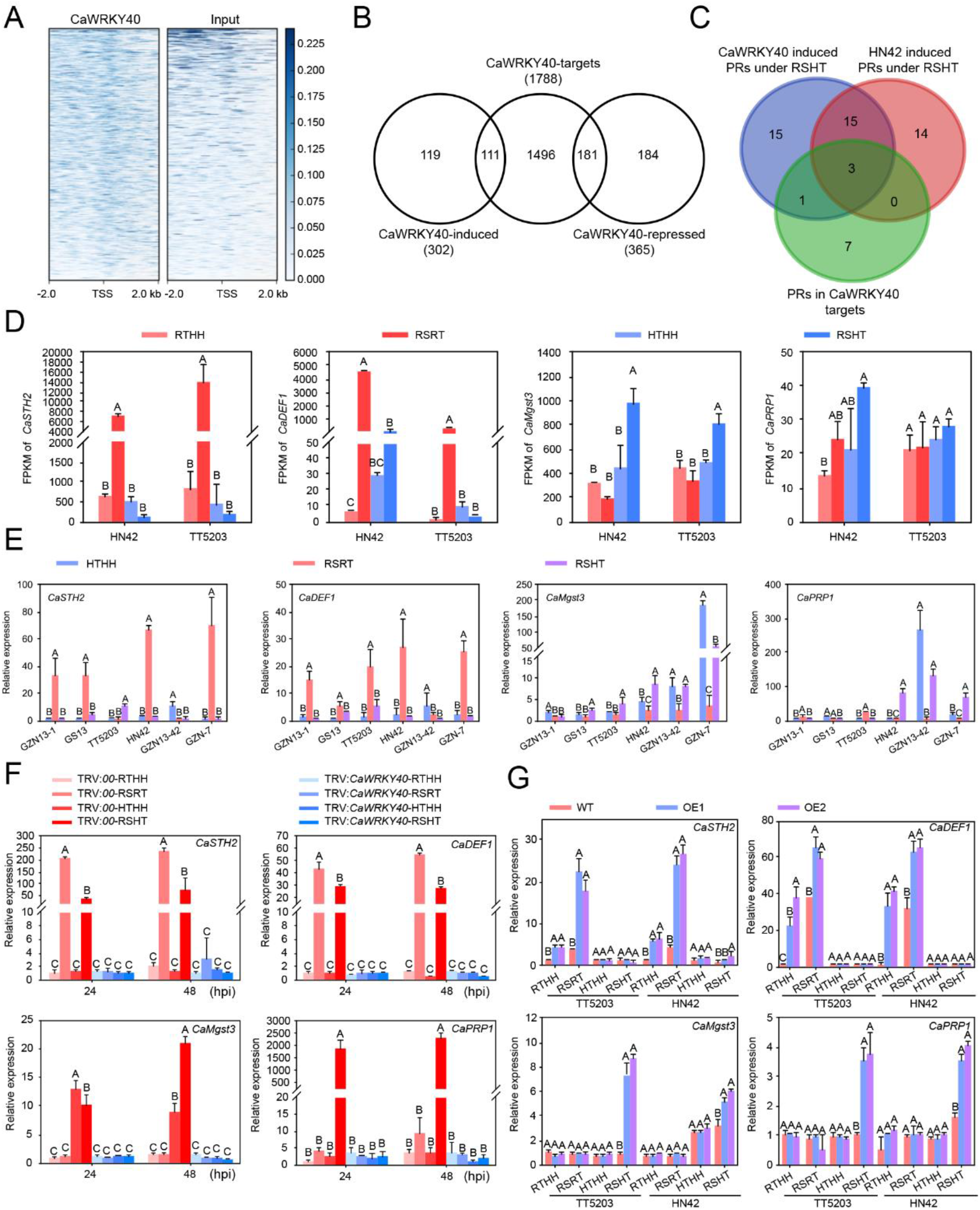
Identification of CaWRKY40-binding Sites and Target Genes upon RSRT or RSHT. (a) Relative binding-peak distribution across CaWRKY40 binding sites near the TSSs. The 2-kbp region upstream of the TSS is defined as the promoter. (b) and (c) Numbers of genes common to CaWRKY40-regulated and -targeted genes as well as CaWRKY40-regulated and -targeted *PR* genes, shown as Venn diagrams (see Table S3 for full results). Roots from HN42 plants, infiltrated with the silencing vectors pTRV:*00* and pTRV:*CaWRKY40*, were challenged with RSHT and harvested for RNA- and ChIP-seq analysis. The association mapping of RNA- and ChIP-seq data were performed to analyze the number of genes upregulated by CaWRKY40. (d) Expression estimates for *STH2, DEF1, Mgst3*, and *PRP1*, as FPKM. (e) Relative transcript levels of *STH2, DEF1, Mgst3*, and *PRP1* in roots of various pepper inbred lines upon RSRT, or RSHT to that upon HTHH, as determined by RT-qPCR. (f) and (g) Relative transcript levels of different *PR* genes in the roots of *CaWRKY40*-silenced HN42 and control plants (f) and TT5203 and HN42 plants overexpressing *CaWRKY40* (g) upon RSRT, HTHH, or RSHT at 24 and 48 hpt to that of *TRV:00* or wild-type plants upon RTHH, as shown by RT-qPCR. In (d), (e), (f), and (g), Data represent the mean ± SD of three biological replicates. Different capital letters above the bars indicate significant difference (*P* < 0.01) by Fisher’s protected LSD test.

To validate *Mgst3, PRP1, STH2*, and *DEF1* as direct targets of CaWRKY40, we performed RT-qPCR. *STH2, DEF1, Mgst3*, and *PRP1* were all downregulated under RSRT by Ca*WRKY40* silencing (Fig. 4f). By contrast, *CaWRKY40* overexpression resulted in upregulation of *Mgst3* and *PRP1* during RSHT, and upregulation of *STH2* and *DEF1* under RSRT (Fig. 4g). CaWRKY40 thus promotes pepper responses to RSRT and RSHT by inducing the expression of such *PR* genes as *STH2* and *DEF1* (in RSRT) and *Mgst3* and *PRP1* (in RSHT).

### CaWRKY40 Differential Targeting Occurs via Selective Binding of W-, WL-, or WT-boxes

To establish how CaWRKY40 regulates distinct target genes under RSRT and RSHT, we determined the consensus sequences within CaWRKY40 binding sites identified by ChIP-seq upon RSRT and RSHT. We identified W-, WL-, and WT-box motifs. To test whether CaWRKY40 indeed binds these motifs, we performed a microscale thermophoresis (MST) assay. We tested CaWRKY40-GST expressed in *Escherichia coli* and CaWRKY40-GFP protein isolated from pepper leaves (Fig. 5b–5c) against synthetic promoter fragments containing 54 bp of a minimal promoter from the cauliflower mosaic virus promoter and three copies of the W-, WL-, or WT-box motifs or mutated versions (designated boxm). *Ca*WRKY40-GST bound DNA fragments containing W-, WL-, or WT-boxes, but not those containing the mutated versions (Fig. 5d).

**Fig. 5.**
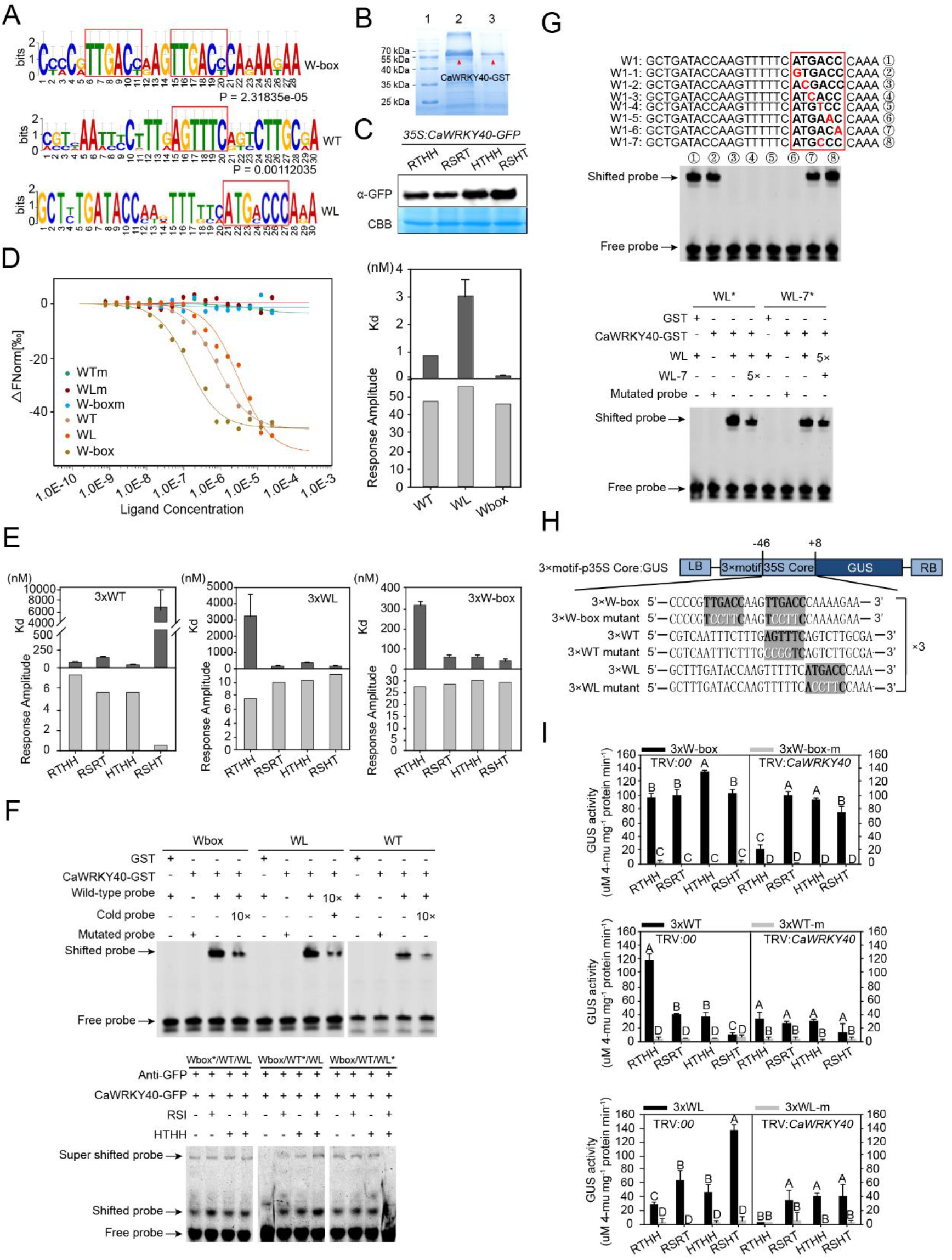
CaWRKY40 Shows Differential Binding to W-, WL- and WT-box *Cis* elements upon RSRT or RSHT. (a) Conserved DNA elements enriched within 500 bp from CaWRKY40 binding-peak regions by MEME, shown as sequence logos. (b) Purification of CaWRKY40-GST expressed in *E. coli*. (c) Detection of CaWRKY40-GFP in roots of pepper plants overexpressing CaWRKY40-GFP upon RTHH, RSRT, HTHH, or RSHT by immunoblot. α-GFP, anti-GFP antibody; CBB, Coomassie Brilliant Blue. (d) Binding of CaWRKY40-GST to W-, WL-, and WT-box determined by MST assay. (e) Binding of CaWRKY40-GFP to W-, WL-, and WT-box determined by MST assay. The CaWRKY40- GFP protein was extracted from the roots of pepper plants overexpressing CaWRKY40-GFP and challenged with RSRT, HTHH, or RSHT and purified by GFP-trap agarose beads. (f) Binding of CaWRKY40-GST or CaWRKY40-GFP to W-, WL-, and WT-box determined by EMSA. (g) Core sequence TGA(C)C of WL-box bound by CaWRKY40-GST determined by EMSA. (h) Schematic map of synthetic promoters containing 3×W-, 3×WL-, and 3×WT-box or their mutants. (i) Activity of GUS driven by synthetic promoters containing 3×W-, 3×WL-, and 3×WT-box or their mutants, in leaves of *CaWRKY40*-silenced or control pepper plants upon RTHH, RSRT, HTHH, or RSHT at 48 hpt. Data represent the mean ± SD of three biological replicates. Different capital letters above the bars indicate significant difference (*P* < 0.01) by Fisher’s protected LSD test.

Based on response amplitude and estimated *K*_d_ values for each motif, CaWRKY40-GFP displayed low binding affinity to the WT-box upon RSHT and low affinity to the WL- and W-box motifs upon RSRT. By contrast, CaWRKY40 associated with the WT-box (upon RSRT) and the WL-box motif (upon RSHT) with high affinity (Fig. 5e). Electrophoresis mobility shift assays (EMSA) with CaWRKY40-GST protein expressed in *E. coli* and Cy5-labeled DNA containing the same synthetic DNA fragments (consisting of the 35S minimal promoter and three copies of W-, WL- or WT-box) revealed clear interaction (mobility shift) between CaWRKY40-GST and W-, WL- or WT-box (Fig. 5f).

We assessed selective interactions between the three boxes and CaWRKY40 in competition experiments by EMSA with pepper plants overexpressing CaWRKY40-GFP and challenged with RSRT or RSHT: excess WT-box probe competed for binding with CaWRKY40-GFP only upon RSRT, whereas excess WL-box probe competed for binding by CaWRKY40-GFP only upon RSHT (Fig. 5f). We scanned the WL-box by testing binding between CaWRKY40-GST and single-base-pair mutated versions of the WL-box: the core sequence TGAC within the WL-box was crucial for CaWRKY40-GST binding to the WL-box. While both WL-box (ATGACC) and point mutant WL-7 (ATGCCC) showed binding by CaWRKY40-GST in EMSA, competition experiments with a 5× excess of WL-box (with 1× WL-7) or a 5× excess of WL-7 (with 1× WL-box) demonstrated efficient competitive binding of each probe by the CaWRKY40-GST protein. These results indicate that CaWRKY40 displays similar binding affinity towards ATGACC and ATGCCC (Fig. 5g).

We studied the effect of CaWRKY40 selective binding on the expression of the *β-GLUCURONIDASE* (*GUS*) gene under the control of synthetic promoter fragments containing the 35S minimal promoter and two copies of the WT-box (or W- or WL-box)., *CaWRKY40* silencing markedly reduced the GUS activity from the WT-box synthetic promoter under RSRT and from the WL-box synthetic promoter under RSHT (Fig. 5h–5i). These results indicate that CaWRKY40 acts as positive regulator upon RSHT by selectively binding the WL-box motif, whereas it selectively targets the WT-box under RSRT.

### CaWRKY40 Differentially Modulates *IPT5, ICS1*, and *PR* Genes via the W-, WL-, and WT-boxes

We selected two other CaWRKY40 target genes, namely *IPT5* and *ICS1*, whose promoters contain only WL-and WT-box motifs, respectively, to assess their possible regulation by CaWRKY40 via these promoter *cis* elements. *IPT5* exhibited higher transcript levels in the roots of control plants and those under RSHT at all time points, although *IPT5* transcript levels remained low upon RSRT or HTHH except at 1 hpt. By contrast, *ICS1* was upregulated by RSRT at all time points, but was upregulated by RSHT only before 3 hpt and downregulated thereafter (Fig. 6a). *ICS1* and *IPT5* were differentially regulated by CaWRKY40: *ICS1* was positively regulated by CaWRKY40 only upon RSRT and displayed decreased transcript levels in the roots of *CaWRKY40*-silenced plants (Fig. 6b) but higher levels in the roots of HN42 and TT5203 plants overexpressing *CaWRKY40* (Fig. 6c) as well as in *N. benthamiana* plants (Fig. 6d). In contrast to *ICS1, IPT5* was positively regulated by CaWRKY40 only upon RSHT, as evidenced by *IPT5* downregulation in *CaWRKY40*-silenced plants upon RSHT and upregulation in pepper plants (Fig. 6c) and *N. benthamiana* plants (Fig. 6d) overexpressing *CaWRKY40*. The upregulation of *ICS1* upon RSHT before 3 hpt was *CaWRKY40*-independent, since *CaWRKY40* silencing did not alter *ICS1* transcript levels (Fig. 6b).

**Fig. 6.**
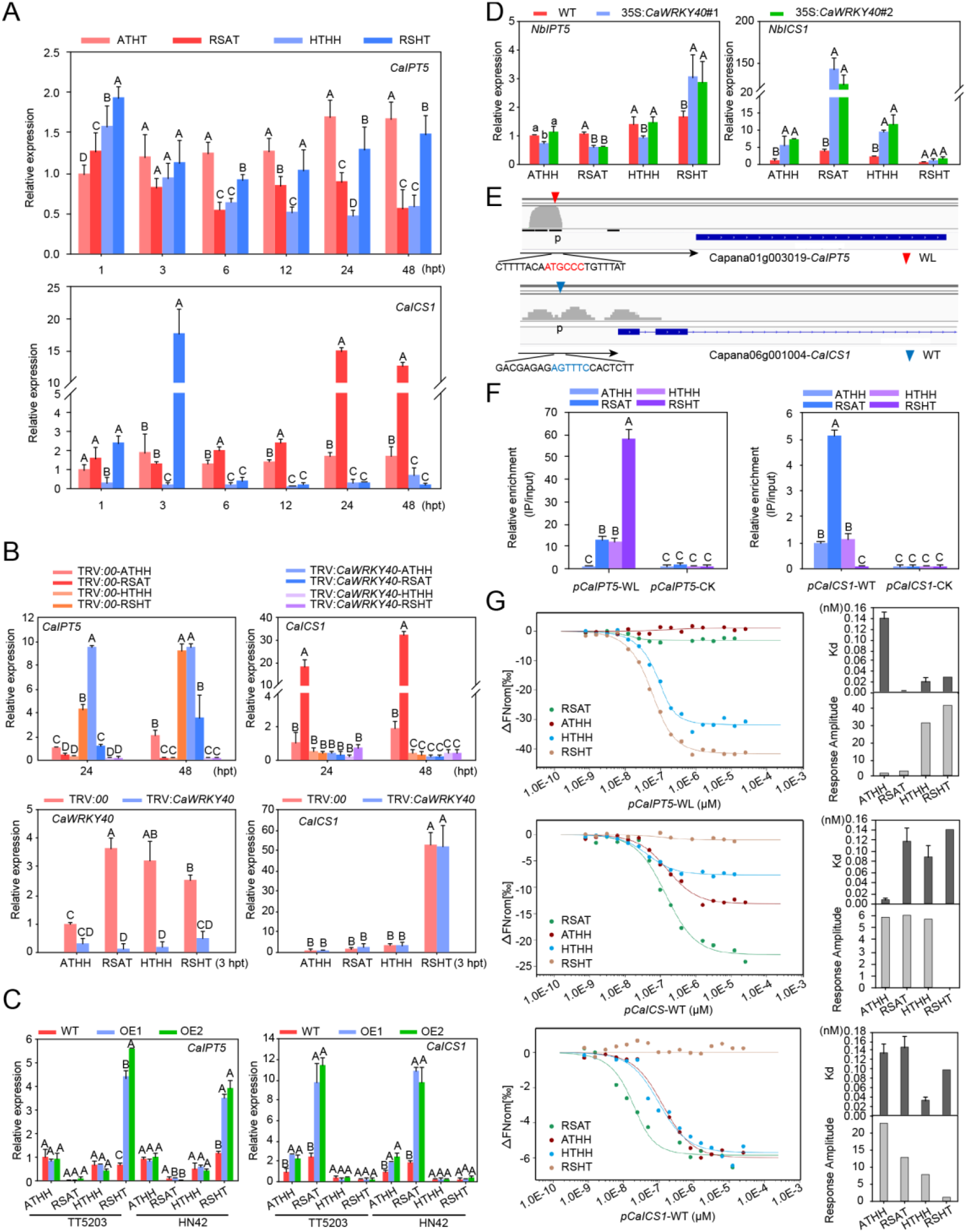
CaWRKY40 Regulates *ICS1* upon RSRT via WT-box and Regulates *IPT5* upon RSHT via WL-box. (a) Transcriptional profiling of *ICS1* and *IPT5* in roots of the pepper inbred line HN42 challenged by RTHH, RSRT, HTHH, or RSHT at different time points. (b) Relative transcript levels of *ICS1* and *IPT5* in roots of *CaWRKY40*-silenced HN42 and control pepper plants upon RSRT, HTHH, or RSHT at 3, 24, and 48 hpt to that of TRV:00 plants upon RTHH, as determined by RT-qPCR. (c) Relative transcript levels of *ICS1* and *IPT5* in roots of HN42 and TT5203 pepper plants overexpressing *CaWRKY40* and their control plants upon RSRT, HTHH, or RSHT at 48 hpt to that of TT5203 wild-type plants upon RTHH. (d) Relative transcript levels of *NbICS1* and *NbIPT5* in roots of *N. benthamiana* plants expressing *CaWRKY40* upon RSRT, HTHH, or RSHT at 48 hpt to that of wild-type plants upon RTHH. (e) Integrative Genomics Viewer (IGV) images of ChIP-seq data and the location of WL- and WT-box within the *ICS1* and *IPT5* promoters. (f) Relative enrichment of CaWRKY40 at the WT- and WL-box within the *ICS1* and *IPT5* promoters in the roots of HN42 plants overexpressing *CaWRKY40*, upon RTHH, RSRT, HTHH, or RSHT at 48 hpt to that upon RTHH, as determined by ChIP-qPCR. (g) Binding of CaWRKY40-GFP to the WL- and WT-box within the *ICS1* and *IPT5* promoters and their corresponding *K*_d_ and response amplitude, as determined by MST. The CaWRKY40-GFP protein was extracted from the roots of HN42 plants overexpressing CaWRKY40-GFP and challenged with RSRT, HTHH, or RSHT at 3 hpi (for g3) and 48 hpi (for g1 and g2) and purified by GFP-trap agarose beads. In (a), (b), (c), (e), and (g), data represent the mean ± SD of three biological replicates. Different capital letters above the bars indicate significant difference (*P* < 0.01) by Fisher’s protected LSD test.

We tested for direct targeting of *ICS1* and *IPT5* by CaWRKY40 and its association with the WL- and WT-boxes within their promoters by ChIP-qPCR. The deposition of CaWRKY40 on the WL-box-containing *IPT5* promoter was enhanced only by RSHT, whereas the amount of CaWRKY40 on the WT-box-containing *ICS1* promoter was specifically enhanced by RSRT (Fig. 6e–6f). We obtained similar results in MST assays using CaWRKY40-GFP expressed in pepper plants under RSRT or RTHT with *IPT5* and *ICS1* promoter fragments containing the WT- or WL-box (Fig. 6g). Collectively, these data indicate that CaWRKY40 positively regulates *ICS1* only upon RSRT via the WT-box, and positively regulates *IPT5* only upon RSHT via the WL-box. In agreement with these observations, expression of *CaWRKY40* in *N. benthamiana* plants led to higher tZ, but not SA, content upon RSHT, and higher SA, but not tZ, content under RSRT (Fig. S5).

We tested CaWRKY40 binding to the *STH2, DEF1, Mgst3*, and *PRP1* promoters by ChIP-PCR using chromatin purified from pepper roots overexpressing Ca*WRKY40-GFP* with specific primers targeting promoter fragments containing the W-,WT-, or WL-box. CaWRKY40 was highly enriched at promoter fragments containing the W-,WT-, or WL-box, but not those lacking these *cis elements* (Fig. S6), indicating that CaWRKY40 may regulate the different sets of *PR* genes via the W-, WL-, and WT-boxes.

### *CaIPT5* and Cytokinin Act Positively in Pepper Responses to RSHT

The increase in *IPT5* transcript and accumulation of tZ in pepper roots under RSHT suggested the involvement of cytokinins in the response to RSHT. To test this hypothesis, we assessed the roles of *IPT5* and tZ in pepper responses to RSHT. Silencing of *IPT5* by VIGS markedly lowered the resistance of pepper plants (Fig. 7c, 7e–7f), while *IPT5* expression enhanced *N. benthamiana* resistance to RSHT (Fig. 7a– 7b, and 7d). Consistent with these results, the specific RSHT-responsive genes *Mgst3* and *PRP1* were significantly downregulated by *IPT5* silencing in pepper plants, and *IPT5* expression in *N. benthamiana* plants induced the expression of their orthologues *NbMgst3* and *NbPRP1* under RSHT. By contrast, the specific RSRT-responsive genes *STH2* and *DEF1* were not downregulated by *CaIPT5* silencing. Likewise, *IPT5* overexpression in *N. benthamiana* did not affect *NbSTH2* and *NbDEF1* expression (Fig. S7a–S7b).

**Fig. 7.**
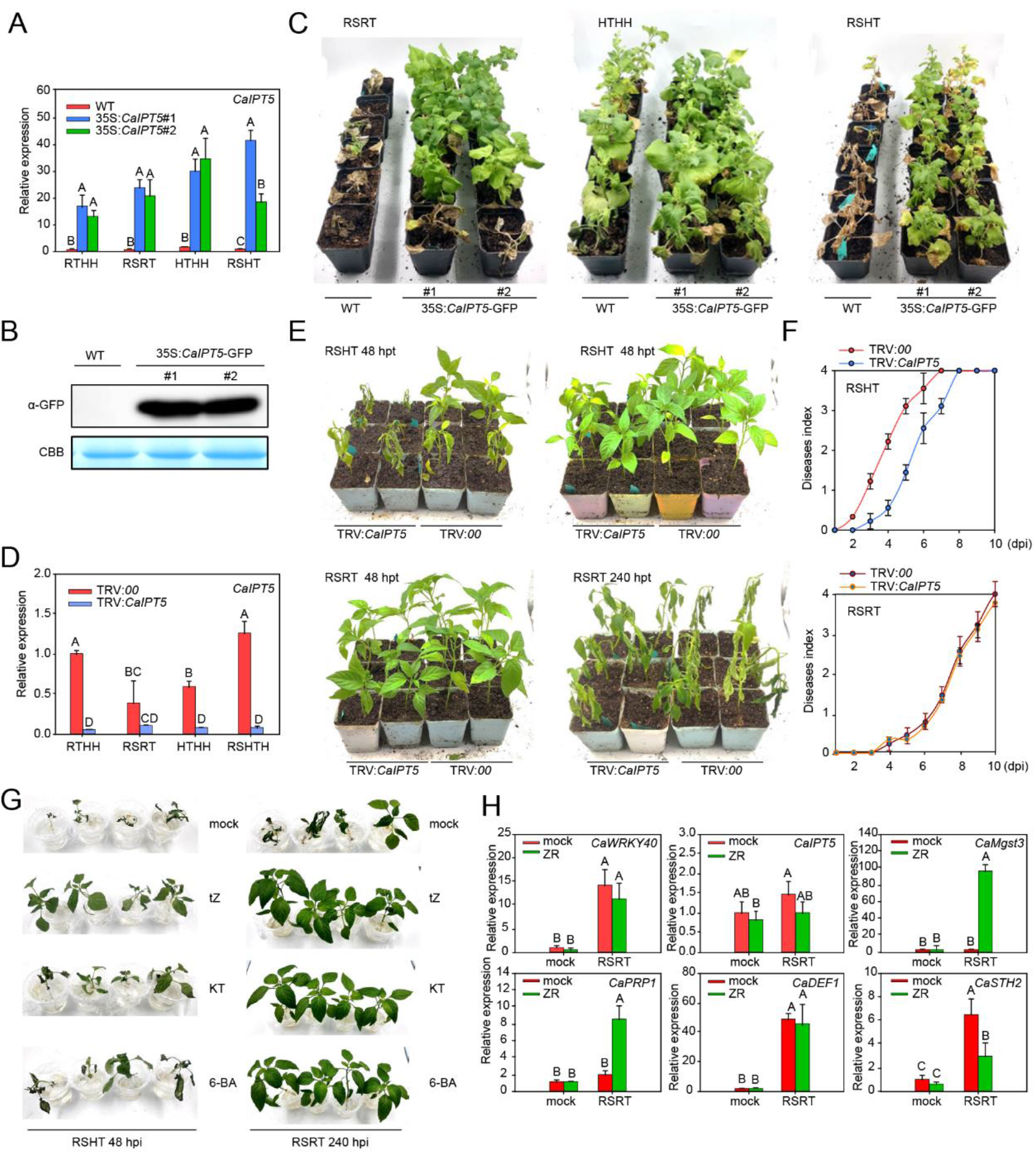
Pepper Immunity to RSHT Is Positively Regulated by CK Signaling. (A, B) Relative transcript levels of *IPT5* by RT-qPCR (a) and IPT5 protein levels by immunoblot (b) in *N. benthamiana* plants expressing *IPT5*. α-GFP, anti-GFP antibodies; CBB, Coomassie Brilliant Blue. (c) Phenotypes of *N. benthamiana* plants expressing *IPT5* and challenged by RSRT, HTHH, or RSHT at 48 hpt. (d) Efficiency of *IPT5* silencing in roots of pTRV:*IPT5* HN42 pepper plants upon RTHH, RSRT, HTHH and RSHT, as seen by the relative expression of *IPT5* to the control by RT-qPCR at 48 hpt. (e) Phenotypic responses of *IPT5-*silenced pepper plants to RSRT or RSHT at 48 hpt and 240 hpt. (f) Disease index of *IPT5*-silenced pepper plants from 0 to 10 dpi. (g) Phenotypes of HN42 pepper plants treated with tZ, KT, or 6-BA (using ddH_2_O as mock treatment) under RSHT at 48 hpt and under RSRT at 240 hpt, respectively. (h) Relative transcript levels of *CaWRKY40, IPT5, Mgst3, PRP1, DEF1*, and *STH2* in HN42 pepper plant roots treated with tZ with or without inoculation with *R. solanacearum* by RT-qPCR at 48 hpt compared to that of the mock. In (a), (d), and (h), data represent the mean ± SD of three biological replicates. Different capital letters above the bars indicate significant difference (*P* < 0.01) by Fisher’s protected LSD test.

In addition, *IPT5* silencing significantly reduced tZ accumulation in the roots of pepper plants (Fig. S8), suggesting that higher tZ levels, resulting from the activation of *IPT5* expression by CaWRKY40, may play a role in pepper responses to RSHT. To test this hypothesis, we analyzed the effect of exogenous tZ application on the response of pepper plants to RSHT. Exogenous tZ significantly enhanced pepper resistance to RSHT, accompanied by upregulation of *Mgst3* and *PRP1* and downregulation of *STH2* and *IPT5*, while *DEF1* and *CaWRKY40* transcript levels did not change (Fig. 7g–7h). Importantly, exogenous tZ also enhanced the resistance of tomato and tobacco plants to RSHT, whereas neither SA nor JA enhanced tolerance to RSHT (from 34 to 37°C, 90% humidity). However, exogenous application of JA or SA did significantly enhance tolerance of tomato and tobacco plants to RSRT (Fig. S9). These results suggest that signaling mediated by CK acts as a positive regulator of tolerance to RSHT in the Solanaceae, including pepper, while plant immunity mediated by SA signaling and JA signaling may not participate in the regulation of Solanaceae responses to RSHT.

### tZ Modifies CaWRKY40 Targeting of *Mgst3, PRP1, IPT5*, and *ICS1* During Pepper Responses to RSHT

As tZ and CaWRKY40 act positively in pepper responses to RSHT, we hypothesized that tZ function might be related to CaWRKY40. To test this hypothesis, we compared the effects of RSHT on chromatin remodeling of *Ca*WRKY40 target genes and the potential role of tZ in this process by measuring the deposition of the histone marks H3K4me3 and H3K9me2, which are correlated with active and inactive chromatin, respectively (Lusser, 2002; Mathieu *et al*., 2005). Upon RSHT, or when treated with tZ and inoculated with *R. solanacearum*, the *Mgst3, PRP1*, and *IPT5* loci exhibited a high H3K4me3 and low H3K9me2 deposition within their transcriptional start sites (TSSs) (Fig. 8a–8b) or promoters (Fig. 8c), while *STH2, DEF1*, and *ICS1* displayed a high H3K4me3 and low H3K9me2 deposition within their TSSs (Fig. 8a–8b) or promoters upon RSRT (Fig. 8c). These results demonstrate that chromatin activation and inactivation are involved in the differential targeting by *Ca*WRKY40 upon RSHT and RSRT.

**Fig. 8.**
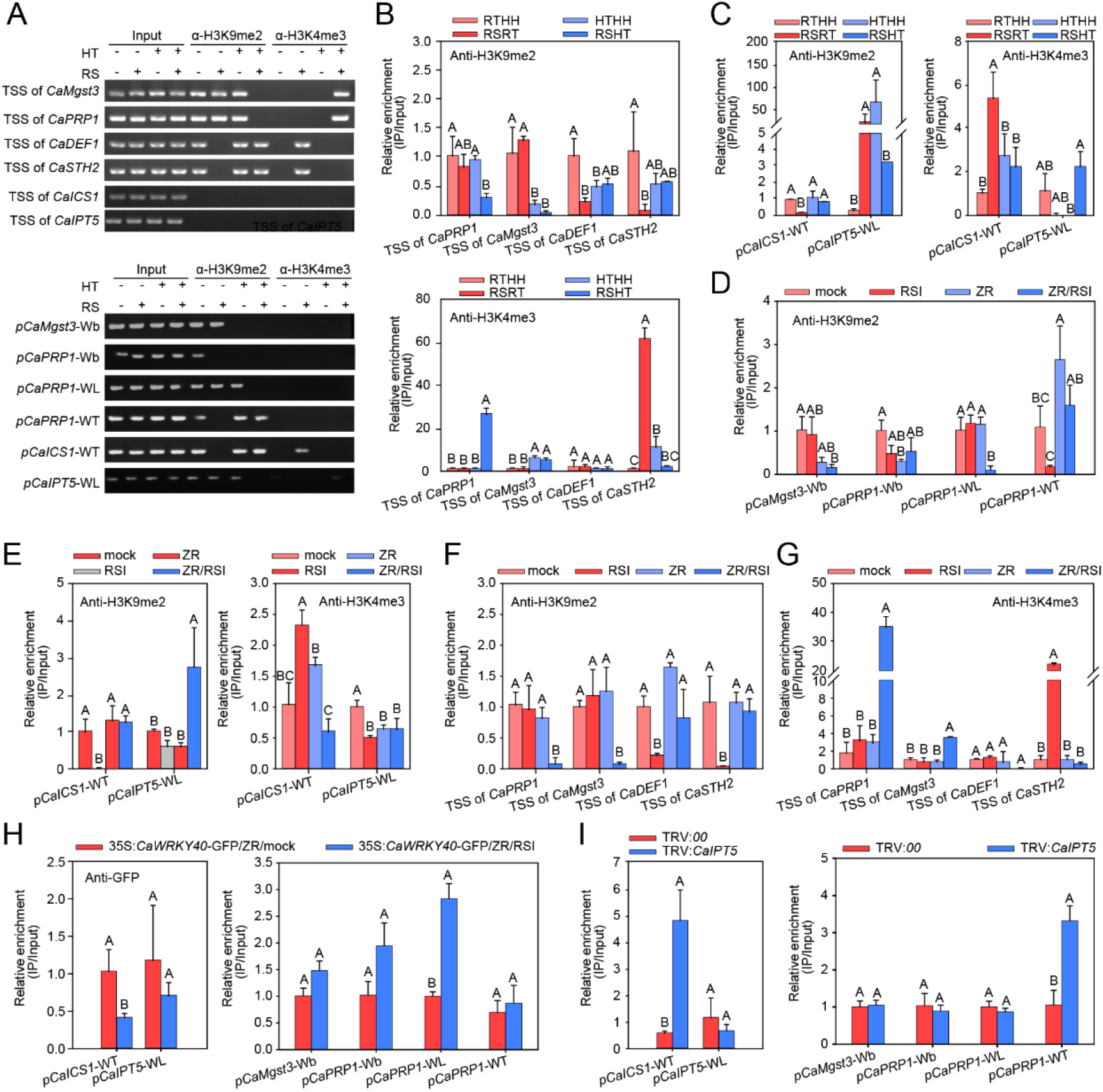
Exogenous Application of tZ Facilitates Targeting of CaWRKY40 to RSHT-specific Responsive *PR* genes. (a) Deposition of H3K9Me2 and H3K4Me3 within the TSSs W-, WL-, or WT-box-containing promoters of different genes upon RTHH, RSRT, HTHH, or RSHT determined by ChIP-PCR. (b) Relative enrichment of H3K9Me2 and H3K4Me3 within the TSSs and promoters of different *PR* genes upon RSRT, HTHH or RSHT to that upon RTHH, determined by ChIP-qPCR. (c) Relative enrichment of H3K9Me2 and H3K4Me3 at the WL- and WT-box within the *ICS1* and *IPT5* promoters upon RSRT, HTHH or RSHT to that upon RTHH, determined by ChIP-qPCR. (d), (e), (f) and (g) Relative enrichment of H3K9Me2 and H3K4Me3 at TSSs and promoters of *Mgst3, PRP1, DEF1, STH2, ICS1*, and *IPT5* upon exogenous application of tZ, with or without inoculation with *R. solanacearum* compared to that of the mock treatment, determined by ChIP-qPCR at 48 hpt. (h) W-, WL- or WT-box dependent binding of CaWRKY40 to the promoters of different genes by ChIP-qPCR in HN42 pepper plant leaves transiently overexpressing *CaWRKY40* upon exogenous application of tZ with or without inoculation with *R. solanacearum* at 48 hpt. (i) W-, WL-, or WT-box dependent binding of CaWRKY40 to the promoters of different genes by ChIP-qPCR in leaves of *IPT5*-silenced pepper plants transiently overexpressing *CaWRKY40* upon exogenous application of tZ with or without inoculation with *R. solanacearum* at 48 hpt. In (a) to (g), the roots of HN42 pepper plants were harvested at 48 hpt for ChIP assay. The enrichment of H3K9Me2 and H3K4Me3 was assessed by PCR or qPCR to amplify the DNA fragments using specific primers (Table S1). In (b) to (i), data represent the mean ± SD of three biological replicates. Different capital letters above the bars indicate significant difference (*P* < 0.01) by Fisher’s protected LSD test.

Exogenous application of tZ to *R. solanacearum*-inoculated plants mimicked RSHT in that it reduced the enrichment of H3K9me2, and enhanced that of H3K4me3, within the TSS of *Mgst3* or *PRP1*, and enhanced H3K9me2 and reduced H3K4me3 deposition to the TSS or promoter of *STH2, DEF1*, and *ICS1* (Fig. 8e–8g). However, unlike RSHT, exogenous application of tZ on plants inoculated with *R. solanacearum* did not decrease enrichment of H3K4me3 at the *IPT5* promoter (Fig. 8c and 8e). These results indicate that pepper plants were primed to activate RSHT-responsive *PR* genes and to inactivate RSRT-responsive *PR* genes upon RSHT via CK signaling. In addition, we assayed the effect of exogenous tZ application to plants inoculated with *R. solanacearum* on the targeting of CaWRKY40 to *PR* genes: CaWRKY40 binding to either *ICS1* or *IPT5* promoters was reduced, but the W-box- or WL-box-dependent binding of CaWRKY40 to the *Mgst3* or *PRP1* promoters was enhanced (Fig. 8h). By contrast, *IPT5* silencing enhanced the WT-dependent binding of CaWRKY40 to the *ICS1* and *PRP1* promoters, but decreased the W- or WL-box-dependent binding of CaWRKY40 to the *ICS1, Mgst3* and *PRP1* promoters (Fig. 7i). These results indicate that the W- or WL-box-dependent binding of CaWRKY40 to its RSHT-responsive target genes is positively regulated, but the WT-box-dependent binding of CaWRKY40 to its target genes is negatively regulated, by CK signaling.

## Discussion

Plants from the Solanaceae family frequently suffer from diseases caused by soil-borne pathogens, including *R. solanacearum*. Although HTHH always worsens the symptoms associated with the diseases (Zhou *et al*., 2004; Koeda *et al*., 2012), how Solanaceae respond to the combination of *R. solonacearum* infection and HTHH (RSHT) remains under-investigated. Here we demonstrate that the pepper plant response to RSHT is distinct from that to RSRT: CaWRKY40 acts synergistically with CK signaling to activate specific immunity to RSHT by transcriptional upregulation of *PR* genes such as *Mgst3* and *PRP1* in a WL-box-dependent manner.

### Pepper Immunity Responses to RSHT and RSRT Are Distinct

Our data indicate that pepper responses to RSRT are largely regulated by immune signaling mediated by the phytohormones SA and JA, with SA signaling activated at an early stage and JA signaling at the later stage. This result is in agreement with the hemibiotrophic lifestyle of the pathogenic bacterium *R. solanacearum*, as well as with the well-established concept that SA and JA act positively in plant immunity against biotrophic and necrotrophic pathogens, respectively (Glazebrook, 2005). Upon RSHT, SA signaling was enhanced in pepper roots early during exposure to RSHT (before 3 hpi) but declined significantly thereafter, while JA signaling was impaired during the entire infection period. However, we show here that CK signaling was activated to lower the susceptibility of pepper plants to RSHT, since exogenous application of tZ or overexpression of *IPT5* significantly raised the resistance of pepper plants to RSHT (Fig. 7). Importantly, immunity mediated by CK signaling was closely linked to the upregulation of GST-encoding genes, including *Mgst3* and *PRP1* (Fig. 6 and 8). A role for *GST* genes in disease resistance is supported by previous genome-wide association studies (GWAS) showing that a GST-encoding gene contributes substantially to wilt resistance in cotton (*Gossypium hirsutum*) (Gong *et al*., 2018). In addition, another GST-encoding gene, *Fusarium head blight 7* (*Fhb7*), was recently shown to confer broad resistance to head blight from *Fusarium* species in wheat (*Triticum aestivum*) (Wang *et al*., 2020), resistance that was improved by moderately high temperatures and high humidity (Dufault *et al*., 2006; Manstretta & Rossi, 2016; Krnjaja *et al*., 2018). It can be concluded that pepper resistance to RSRT is positively regulated by SA and JA signaling, but that its resistance to RSHT is regulated by CK signaling. This hypothesis is supported by our observations that the resistance of pepper, tomato, and tobacco plants to RSRT was enhanced by exogenously applied SA or JA. By contrast, their resistance to RSHT was not significantly affected by exogenously applied SA or JA, although it was ameliorated by exogenous application of tZ (Fig. S9), indicating that this mechanism is conserved across Solanaceae plants.

### CaWRKY40 Acts Positively in Pepper Resistance to RSHT and RSRT by Differentially Targeting *PR* Genes

The data collected from our loss- and gain-of-function assays demonstrate that CaWRKY40 acts positively in pepper responses to both RSRT and RSHT. Importantly, the results from our comprehensive analysis of RNA-seq and ChIP-seq datasets linked the function of CaWRKY40 in RSHT resistance with target genes such as *IPT5* and the *PR* genes *Mgst3* and *PRP1*, while providing a link between the function of CaWRKY40 in RSRT resistance and the target gene *ICS1* and the *PR* genes *STH2* and *DEF1*. Among these target genes, *IPT5* phenocopied *CaWRKY40* in terms of resistance to RSHT treatment when silenced or overexpressed, and exerted effects similar to CaWRKY40 on the expression of *Mgst3* and *PRP1* (Fig. 3–5). In agreement with this, the overexpression of *CaWRKY40* upon RSHT enhanced accumulation of tZ (Fig. S5), whose exogenous application significantly enhanced transcript levels of *Mgst3* and *PRP1*, to a degree similar to that seen upon overexpression of *CaWRKY40* or *IPT5* (Fig. 4g, S7b, and 6h). These results indicate that CK signaling is crucial for pepper resistance to RSHT that is activated by CaWRKY40, and that CaWRKY40 may regulate pepper resistance to RSHT through the upregulation of *IPT5* expression, and therefore CK signaling. Notably, when tZ was used alone, neither *Mgst3* or *PRP1* expression could be induced, similar to the result of a previous study in which CK signaling was found to exert its function in plant immunity only in the presence of pathogens or elicitors (Conrath *et al*., 2015). We speculate that, as CKs are crucial regulators in plant growth and development, their signaling is tightly regulated to avoid automatically activating plant immunity in the absence of pathogens.

### The Specific Targeting of CaWRKY40 in Pepper Responses to RSHT Is Regulated by CK Signaling via Coordinated Chromatin Activation and WL-box Binding

CaWRKY40 fulfills its function in positively regulating pepper resistance to RSHT and RSRT through differential gene targeting, which might be determined at first by the chromatin state (Garner *et al*., 2016). We determined that the application of exogenous tZ to plants inoculated with *R. solanacearum* mimicked RSHT in causing chromatin activation of the *Mgst3* or *PRP1* loci, as they displayed reduced enrichment of the chromatin inactivation mark H3K9me2 (Mathieu *et al*., 2005) and increased enrichment of the chromatin activation mark H3K4me2 (van Dijk *et al*., 2005). By contrast, chromatin inactivation of *ICS1, STH2*, or *DEF1* manifested itself by the enrichment of H3K9me2 or the reduced deposition of H3K4me2 at their TSSs, which were conferred by RSHT and by exogenously applied tZ on plants inoculated with *R. solanacearum* (Fig. 8). These data indicate that, upon RSHT, CK signaling confers chromatin activation of RSHT-responsive and CaWRKY40-target genes, while also inactivating the chromatin around RSRT-responsive and CaWRKY40-target genes.

Another key step in the targeting of a given TF is its selective binding to its cognate *cis*-element within the promoters of target genes. Within the three potential cognate Ca*WRKY40 cis elements* identified by ChIP-seq, the W-box is bound by the majority of WRKY TFs that have been functionally characterized (Rushton *et al*., 2010). The WT-box is selectively bound by WRKY26, WRKY33, WRKY41, or WRKY70 in *Arabidopsis thaliana* (Machens *et al*., 2014; Liu *et al*., 2015; Kanofsky *et al*., 2017; Kanofsky *et al*., 2018). The novel WL-box motif identified here has not been previously investigated. We provide strong evidence that the WL-box is bound by CaWRKY40 upon RSHT, with the core sequence TGA(C)C in the WL-box being crucial for this binding (Fig. 5f–5i). Importantly, we observed that in plants inoculated with *R. solanacearum*, either RSHT or exogenous tZ application enhanced the binding of CaWRKY40 to the W- or WL-box, while reducing its binding to the WT-box. The selective binding of CaWRKY40 to the three different *cis elements* under different conditions might be attributable to some as yet unidentified signaling component(s) differentially activated by these various stresses, since WRKY TFs may interact with other proteins that will modulate their targeting or functions (Chi *et al*., 2013).

The regulation of *IPT* genes by CaWRKY40 appears to be comparatively complex. Our RNA-seq data showed that *IPT2, IPT5*, and *IPT9* in pepper were all upregulated by RSHT, but the putative Arabidopsis orthologues *IPT2* and *IPT9* have no transferase activity and may therefore not be involved in CK biosynthesis (Takei *et al*., 2001). Thus, we focused on *IPT5*, which was positively regulated by CaWRKY40 in pepper roots upon RSHT, but was negatively regulated by CaWRKY40 upon RTHH, as the low transcript levels of Ca*WRKY40* in non-stressed pepper roots or Ca*WRKY40*-silenced pepper plants resulted in higher *IPT5* expression (Fig. 6). However, this upregulation of *IPT5* did not result in a rise in CK content, including tZ, which might be due to repression by high levels of indoleacetic acid (IAA) (Bielach *et al*., 2012). Besides the activation of *IPT5* by CaWRKY40, the higher tZ content in the roots of pepper plants challenged by RSHT might also be partially due to the derepressive effect of IAA on CK biosynthesis, as the IAA content in the roots of RSHT-challenged pepper plants was much lower than that of control plants (Fig. 2c). Our data also indicate that CK biosynthesis might be subject to negative feedback regulation when the concentration of CKs such as tZ is too high, since exogenously applied tZ, unlike RSHT that induced *IPT5* expression, converted the chromatin around the *IPT5* locus to a more repressed state by increasing H3K9me2 and reducing H3K4me2 deposition to its TSS or promoter, and also blocked the binding of CaWRKY40 to the WL-box within the IPT5 promoter (Fig. 8e and 8h).

Collectively, the results of the present study show that pepper immunity in response to *R. solanacearum* infections under HTHH (RSHT) is distinct from that to *R. solanacearum* infections at ambient temperature (RSRT). Upon RSHT, upregulated CaWRKY40 positively regulates *IPT5*, and the resulting increase in tZ content in turn synergistically works with CaWRKY40 to activate specific immunity to RSHT by coordinately promoting chromatin activation and promoter binding via the WL-box, while disabling CaWRKY40-mediated activation of RSRT immunity by inactivating the chromatin of specific RSRT-responsive genes and repressing binding to promoters containing the WT-box.

## Supporting information

supplementary-informatio

## ACKNOWLEDGMENTS

The authors thank Mark D. Curtis for kindly providing the Gateway destination vectors and Dr. S. P. Dinesh-Kumar of Yale University for the pTRV1 and pTRV2 vectors. This work was supported by grants from the National Natural Science Foundation of China (31572136, 31372061), the Development Fund Project of Fujian Agriculture and Forestry University (CXZX2016158, CXZX2017548), and the Scientific Research Foundation of the Graduate School of Fujian Agriculture and Forestry University (324-1122yb047).

## Data Availability Statement

The data that support the findings of this study are openly available in [https://db.cngb.org/cnsa/] at [China National GeneBank], reference number [CNP0001155; CNP0001156; CNP0001157].

## Author contributions

S.L.H and S.Y conceived the research and designed the experiments. S.Y., W.W.C, L.S., R.J.W., J.S.C., J.S.C., S.C.H., Y.T.Z. Q.X.Z. and A.W.W. performed the experiments. S.Y., W.W.C, L.S. and D.Y.G. analyzed the data. S.L.H. wrote the manuscript.

## ABBREVIATIONS

(RT or 28°C): room temperature
(HT or 37°C): high temperature
(HH): high humidity
(RS or RSI): *R. solanacearum* infection
(RSRT): *R. solanacearum* infection at room temperature
(RSHT): *R. solanacearum* infection at high temperature and high humidity
(RTHH): growth at room temperature and high humidity
(HTHH): growth at high temperatures and high humidity

## SUPPLEMENTARY-INFORMATION

Methods

Methods S1. Vector Construction.

Methods S2. Genetic Transformation of Pepper and *N. Benthamiana*.

Methods S3. Virus-Induced Gene Silencing (VIGS)

Methods S4. Exogenous Applications of SA, JA, tZ, KT, or 6-BA

Methods S5. RNA-seq Analysis

Methods S6. EMSA Analysis

Methods S7. MST Analysis

Methods S8. GUS Activity Assay

Methods S9. prokaryotic Expression and Protein Purification

Methods S10. plant Total Protein Extraction and Immunoblot Analysis

Methods S11. phytohormone Measurements and Quantification

## KEY RESOURCES TABLE

### Tables

Table S1. Primers used in this study.

Table S2. Disease index for pepper plants infected with *R. solanacearum*.

Table S3. Venn diagram results for Figure 4c.

### Figures

Fig. S1. Phenotypic Response of Plants from Different Inbred Pepper Lines to RSRT, HTHH, or RSHT Treatment.

Fig. S2. Phenotypic Response of *N. benthamiana* and Tomato (*Solanum lycopersicum*) Plants to RSRT, HTHH, or RSHT Treatment.

Fig. S3. Heatmap Representation of the Transcript Levels of Putative R Genes upon RTHH, RSRT, and RSHT Treatment in the TT5203 and HN42 Pepper Inbred Lines.

Fig. S4. Transcript Levels of Members of the Pepper *WRKY* Gene Family upon RTHH, RSRT, HTHH, or RSHT Treatment.

Fig. S5. The Contents of the Phytohormones SA, ZT, tZ, ABA, and JA-Ile in Roots of *N. benthamiana* Plants Overexpressing *CaWRKY40* and Their Control Plants after Challenge with RSRT or RSHT Treatment at 48 hpt.

Fig. S6. Binding of CaWRKY40 to the *STH2, DEF1, Mgst3*, and *PRP1* Promoters by ChIP-PCR Analysis.

Fig. S7. IPT5-mediated Immunity Participates in Pepper Responses to RSHT but not to RSRT.

Fig. S8. The Phytohormone Content for ABA, JA-Ile, SA, tZ, and ZT in Roots of *CaIPT5*-silenced HN42 and Control Pepper Plants Challenged with RTHH, RSRT, HTHH, or RSHT Treatment at 48 hpt.

Fig. S9. The Effect of Exogenous Application of the Phytohormones SA, MeJA, and tZ on the Response of Pepper, Tomato and Tobacco Plants to RSRT and RSHT Treatments.

## References

Albrecht T, Argueso CT. 2017. Should I fight or should I grow now? The role of cytokinins in plant growth and immunity and in the growth-defence trade-off. Ann Bot 119(5): 725–735.

Andaya VC, Tai TH. 2006. Fine mapping of the qCTS12 locus, a major QTL for seedling cold tolerance in rice. Theor Appl Genet 113(3): 467–475.

Atkinson NJ, Urwin PE. 2012. The interaction of plant biotic and abiotic stresses: from genes to the field. J Exp Bot 63(10): 3523–3543.

Attaran E, Zeier TE, Griebel T, Zeier J. 2009. Methyl salicylate production and jasmonate signaling are not essential for systemic acquired resistance in Arabidopsis. Plant Cell 21(3): 954–971.

Bielach A, Duclercq J, Marhavy P, Benkova E. 2012. Genetic approach towards the identification of auxin-cytokinin crosstalk components involved in root development. Philos Trans R Soc Lond B Biol Sci 367(1595): 1469–1478.

Bieri S, Mauch S, Shen QH, Peart J, Devoto A, Casais C, Ceron F, Schulze S, Steinbiss HH, Shirasu K, et al. 2004. RAR1 positively controls steady state levels of barley MLA resistance proteins and enables sufficient MLA6 accumulation for effective resistance. Plant Cell 16(12): 3480–3495.

Bradford MM. 1976. A Rapid and Sensitive Method for the Quantitation of Microgram Quantities of Protein Utilizing the Principle of Protein-Dye Binding. Analytical Biochemistry 72(1-2): 248–254.

Cai H, Yang S, Yan Y, Xiao Z, Cheng J, Wu J, Qiu A, Lai Y, Mou S, Guan D, et al. 2015. CaWRKY6 transcriptionally activates CaWRKY40, regulates Ralstonia solanacearum resistance, and confers high-temperature and high-humidity tolerance in pepper. J Exp Bot 66(11): 3163–3174.

Chang X, Seo M, Takebayashi Y, Kamiya Y, Riemann M, Nick P. 2017. Jasmonates are induced by the PAMP flg22 but not the cell death-inducing elicitor Harpin in Vitis rupestris. Protoplasma 254(1): 271–283.

Chi Y, Yang Y, Zhou Y, Zhou J, Fan B, Yu JQ, Chen Z. 2013. Protein-protein interactions in the regulation of WRKY transcription factors. Mol Plant 6(2): 287–300.

Ciolkowski I, Wanke D, Birkenbihl RP, Somssich IE. 2008. Studies on DNA-binding selectivity of WRKY transcription factors lend structural clues into WRKY-domain function. Plant Mol Biol 68(1-2): 81–92.

Conrath U, Beckers GJ, Langenbach CJ, Jaskiewicz MR. 2015. Priming for enhanced defense. Annu Rev Phytopathol 53: 97–119.

Cutler AJ, Krochko JE. 1999. Formation and breakdown of ABA. Trends Plant Sci 4(12): 472–478.

Dang FF, Wang YN, Yu L, Eulgem T, Lai Y, Liu ZQ, Wang X, Qiu AL, Zhang TX, Lin J, et al. 2013. CaWRKY40, a WRKY protein of pepper, plays an important role in the regulation of tolerance to heat stress and resistance to Ralstonia solanacearum infection. Plant Cell Environ 36(4): 757–774.

Digonnet C, Martinez Y, Denance N, Chasseray M, Dabos P, Ranocha P, Marco Y, Jauneau A, Goffner D. 2012. Deciphering the route of Ralstonia solanacearum colonization in Arabidopsis thaliana roots during a compatible interaction: focus at the plant cell wall. Planta 236(5): 1419–1431.

Dufault NS, De Wolf ED, Lipps PE, Madden LV. 2006. Role of Temperature and Moisture in the Production and Maturation of Gibberella zeae Perithecia. Plant Dis 90(5): 637–644.

Eulgem T, Somssich IE. 2007. Networks of WRKY transcription factors in defense signaling. Curr Opin Plant Biol 10(4): 366–371.

F R, A dCM, ML dCM, H S, D M, V H, H W, H K. 1992. Coat protein mediated resistance to Plum Pox Virus in Nicotiana clevelandii and N. benthamiana. Plant cell reports 11(1): 30–33.

Garner CM, Kim SH, Spears BJ, Gassmann W. 2016. Express yourself: Transcriptional regulation of plant innate immunity. Semin Cell Dev Biol 56: 150–162.

Glazebrook J. 2005. Contrasting mechanisms of defense against biotrophic and necrotrophic pathogens. Annu Rev Phytopathol 43: 205–227.

Gong Q, Yang Z, Chen E, Sun G, He S, Butt HI, Zhang C, Zhang X, Du X, Li F. 2018. A Phi-Class Glutathione S-Transferase Gene for Verticillium Wilt Resistance in Gossypium arboreum Identified in a Genome-Wide Association Study. Plant Cell Physiol 59(2): 275–289.

Grosskinsky DK, Edelsbrunner K, Pfeifhofer H, van der Graaff E, Roitsch T. 2013. Cis- and trans-zeatin differentially modulate plant immunity. Plant Signal Behav 8(7): e24798.

H L, R D. 2009. Fast and accurate short read alignment with Burrows-Wheeler transform. Bioinformatics (Oxford, England) 25(14): 1754–1760.

Hwang CF, Bhakta AV, Truesdell GM, Pudlo WM, Williamson VM. 2000. Evidence for a role of the N terminus and leucine-rich repeat region of the Mi gene product in regulation of localized cell death. Plant Cell 12(8): 1319–1329.

Jambunathan N, Siani JM, McNellis TW. 2001. A humidity-sensitive Arabidopsis copine mutant exhibits precocious cell death and increased disease resistance. Plant Cell 13(10): 2225–2240.

Jefferson RA. 1987. GUS fusion : β-glucuronidase as a sensitive and versatile gene fusion marker in higher plants. Embo Journal 6.

Jones JD, Dangl JL. 2006. The plant immune system. Nature 444(7117): 323–329.

Kanofsky K, Bahlmann AK, Hehl R, Dong DX. 2017. Combinatorial requirement of W- and WT-boxes in microbe-associated molecular pattern-responsive synthetic promoters. Plant Cell Rep 36(6): 971–986.

Kanofsky K, Strauch CJ, Sandmann A, Moller A, Hehl R. 2018. Transcription factors involved in basal immunity in mammals and plants interact with the same MAMP-responsive cis-sequence from Arabidopsis thaliana. Plant Mol Biol 98(6): 565–578.

Khan MI, Zhang Y, Liu Z, Hu J, Liu C, Yang S, Hussain A, Furqan Ashraf M, Noman A, Shen L, et al. 2018. CaWRKY40b in pepper acts as a negative regulator in response to Ralstonia solanacearum by directly modulating defense genes including CaWRKY40. International Journal of Molecular Sciences 19(5).

Koeda S, Hosokawa M, Kang BC, Tanaka C, Choi D, Sano S, Shiina T, Doi M, Yazawa S. 2012. Defense response of a pepper cultivar cv. Sy-2 is induced at temperatures below 24 degrees C. J Plant Res 125(1): 137–145.

Krnjaja V, Stankovic S, Obradovic A, Petrovic T, Mandic V, Bijelic Z, Bozic M. 2018. Trichothecene Genotypes of Fusarium graminearum Populations Isolated from Winter Wheat Crops in Serbia. Toxins (Basel) 10(11).

Kuc J. 1990. Compounds from plants that regulate or participate in disease resistance. Ciba Found Symp 154: 213–224; discussion 224-218.

Liu H, Dong S, Gu F, Liu W, Yang G, Huang M, Xiao W, Liu Y, Guo T, Wang H, et al. 2017. NBS-LRR Protein Pik-H4 Interacts with OsBIHD1 to Balance Rice Blast Resistance and Growth by Coordinating Ethylene-Brassinosteroid Pathway. Front Plant Sci 8: 127.

Liu S, Kracher B, Ziegler J, Birkenbihl RP, Somssich IE. 2015. Negative regulation of ABA signaling by WRKY33 is critical for Arabidopsis immunity towards Botrytis cinerea 2100. Elife 4: e07295.

Lusser A. 2002. Acetylated, methylated, remodeled: chromatin states for gene regulation. Curr Opin Plant Biol 5(5): 437–443.

M S-D, H D, K T, P B. 2010. PeakAnalyzer: genome-wide annotation of chromatin binding and modification loci. BMC bioinformatics 11: 415.

Machens F, Becker M, Umrath F, Hehl R. 2014. Identification of a novel type of WRKY transcription factor binding site in elicitor-responsive cis-sequences from Arabidopsis thaliana. Plant Mol Biol 84(4-5): 371–385.

Mansfield J, Genin S, Magori S, Citovsky V, Sriariyanum M, Ronald P, Dow M, Verdier V, Beer SV, Machado MA, et al. 2012. Top 10 plant pathogenic bacteria in molecular plant pathology. Mol Plant Pathol 13(6): 614–629.

Manstretta V, Rossi V. 2016. Effects of Temperature and Moisture on Development of Fusarium graminearum Perithecia in Maize Stalk Residues. Appl Environ Microbiol 82(1): 184–191.

Mathieu O, Probst AV, Paszkowski J. 2005. Distinct regulation of histone H3 methylation at lysines 27 and 9 by CpG methylation in Arabidopsis. EMBO J 24(15): 2783–2791.

Meng X, Xu J, He Y, Yang KY, Mordorski B, Liu Y, Zhang S. 2013. Phosphorylation of an ERF transcription factor by Arabidopsis MPK3/MPK6 regulates plant defense gene induction and fungal resistance. Plant Cell 25(3): 1126–1142.

N B, F H, MA S, L P. 1994. Protection against virus infection in tobacco plants expressing the coat protein of grapevine fanleaf nepovirus. Plant cell reports 13(6): 357–360.

Naseem M, Wolfling M, Dandekar T. 2014. Cytokinins for immunity beyond growth, galls and green islands. Trends Plant Sci 19(8): 481–484.

Noutoshi Y, Ito T, Seki M, Nakashita H, Yoshida S, Marco Y, Shirasu K, Shinozaki K. 2005. A single amino acid insertion in the WRKY domain of the Arabidopsis TIR-NBS-LRR-WRKY-type disease resistance protein SLH1 (sensitive to low humidity 1) causes activation of defense responses and hypersensitive cell death. Plant J 43(6): 873–888.

O’Donnell PJ, Schmelz E, Block A, Miersch O, Wasternack C, Jones JB, Klee HJ. 2003. Multiple hormones act sequentially to mediate a susceptible tomato pathogen defense response. Plant Physiol 133(3): 1181–1189.

P M, TL B. 2011. MEME-ChIP: motif analysis of large DNA datasets. Bioinformatics (Oxford, England) 27(12): 1696–1697.

Pandey SP, Somssich IE. 2009. The role of WRKY transcription factors in plant immunity. Plant Physiol 150(4): 1648–1655.

Qiu A, Lei Y, Yang S, Wu J, Li J, Bao B, Cai Y, Wang S, Lin J, Wang Y. 2018. CaC3H14 encoding a tandem CCCH zinc finger protein is directly targeted by CaWRKY40 and positively regulates the response of pepper to inoculation by Ralstonia solanacearum. Molecular plant pathology 19(10): 2221.

Reusche M, Klaskova J, Thole K, Truskina J, Novak O, Janz D, Strnad M, Spichal L, Lipka V, Teichmann T. 2013. Stabilization of cytokinin levels enhances Arabidopsis resistance against Verticillium longisporum. Mol Plant Microbe Interact 26(8): 850–860.

Robert-Seilaniantz A, Grant M, Jones JD. 2011. Hormone crosstalk in plant disease and defense: more than just jasmonate-salicylate antagonism. Annu Rev Phytopathol 49: 317–343.

Rushton PJ, Somssich IE, Ringler P, Shen QJ. 2010. WRKY transcription factors. Trends Plant Sci 15(5): 247–258.

Sakakibara H. 2005. Cytokinin biosynthesis and regulation. Vitam Horm 72: 271–287.

Silva MS, Arraes FBM, Campos MA, Grossi-de-Sa M, Fernandez D, Candido ES, Cardoso MH, Franco OL, MF Grossi-de-Sa. 2018. Review: Potential biotechnological assets related to plant immunity modulation applicable in engineering disease-resistant crops. Plant Sci 270: 72–84.

Spoel SH, Koornneef A, Claessens SM, Korzelius JP, Van Pelt JA, Mueller MJ, Buchala AJ, Metraux JP, Brown R, Kazan K, et al. 2003. NPR1 modulates cross-talk between salicylate- and jasmonate-dependent defense pathways through a novel function in the cytosol. Plant Cell 15(3): 760–770.

Su J, Zhang M, Zhang L, Sun T, Liu Y, Lukowitz W, Xu J, Zhang S. 2017. Regulation of Stomatal Immunity by Interdependent Functions of a Pathogen-Responsive MPK3/MPK6 Cascade and Abscisic Acid. Plant Cell 29(3): 526–542.

Takei K, Sakakibara H, Sugiyama T. 2001. Identification of genes encoding adenylate isopentenyltransferase, a cytokinin biosynthesis enzyme, in Arabidopsis thaliana. J Biol Chem 276(28): 26405–26410.

Thomma BP, Eggermont K, Penninckx IA, Mauch-Mani B, Vogelsang R, Cammue BP, Broekaert WF. 1998. Separate jasmonate-dependent and salicylate-dependent defense-response pathways in Arabidopsis are essential for resistance to distinct microbial pathogens. Proc Natl Acad Sci U S A 95(25): 15107–15111.

Tsuda K, Somssich IE. 2015. Transcriptional networks in plant immunity. New Phytol 206(3): 932–947.

van Dijk K, Marley KE, Jeong BR, Xu J, Hesson J, Cerny RL, Waterborg JH, Cerutti H. 2005. Monomethyl histone H3 lysine 4 as an epigenetic mark for silenced euchromatin in Chlamydomonas. Plant Cell 17(9): 2439–2453.

Walters DR, McRoberts N. 2006. Plants and biotrophs: a pivotal role for cytokinins? Trends Plant Sci 11(12): 581–586.

Wang H, Sun S, Ge W, Zhao L, Hou B, Wang K, Lyu Z, Chen L, Xu S, Guo J, et al. 2020. Horizontal gene transfer of Fhb7 from fungus underlies Fusarium head blight resistance in wheat. Science.

Wasilewska A, Vlad F, Sirichandra C, Redko Y, Jammes F, Valon C, Frei dit Frey N, Leung J. 2008. An update on abscisic acid signaling in plants and more. Mol Plant 1(2): 198–217.

Xiao S, Brown S, Patrick E, Brearley C, Turner JG. 2003. Enhanced transcription of the Arabidopsis disease resistance genes RPW8.1 and RPW8.2 via a salicylic acid-dependent amplification circuit is required for hypersensitive cell death. Plant Cell 15(1): 33–45.

Y Z, T L, CA M, J E, DS J, BE B, C N, RM M, M B, W L, et al. 2008. Model-based analysis of ChIP-Seq (MACS). Genome biology 9(9): R137.

Zhou F, Menke FL, Yoshioka K, Moder W, Shirano Y, Klessig DF. 2004. High humidity suppresses ssi4-mediated cell death and disease resistance upstream of MAP kinase activation, H2O2 production and defense gene expression. Plant J 39(6): 920–932.

