## supplementary-informatio for "Environment-Dependent Switch between Two Immune Systems against *Ralstonia solanacearum* Infections in Pepper"

### Supporting information for materials and methods

#### Methods S1. Vector Construction

All vectors generated in the present study were constructed via Gateway cloning techniques (Invitrogen, Carlsbad, CA, USA) using a series of Gateway-cloning-compatible vectors: pTRV2 for gene silencing, pEarlyGate103 for overexpression fused with GFP, pDEST17 for prokaryotic expression in *E. coli*, pMDC163 for promoter activity assay. All DNA fragments were produced by DNA synthesis and then PCR-amplified with Gateway-cloning-compatible primer pairs (Table S1). We synthesized the full length open reading frames (ORF) with or without termination codon (for overexpression), specific fragments within the coding sequence (CDS) or 3' untranslated regions (UTR) (confirmed by BLAST search against the pepper genome sequence) (for silencing), or synthetic promoters containing different DNA motifs (3×W-box, 3×WT, 3×WL or their mutant versions) (for promoter analysis). All resulting PCR products were cloned into entry vector pDONR207 by BP reaction, and then cloned into various destination vectors by LR reaction after confirmation by sequencing.

#### Methods S2. Genetic Transformation of Pepper and *N. benthamiana*

We introduced the *35Spro:CaWRKY40-GFP* binary vector into *Agrobacterium tumefaciens* strain GV3101. *Agrobacterium* cultures were inoculated into LB medium and grown for 48 h at 28ºC, then collected and resuspended in liquid MS medium at an adjusted $OD_{600}$ = 0.4, supplemented with the surfactant Silwet L-77. We excised one cotyledon and wounded the apical meristem of pepper seedlings with a razor blade (at the 2-cotyledon stage). The wounded seedlings were then used as explants for transformation: they were soaked in the *Agrobacterium* cell suspension for 15 min, cleaned and patted dry on clean filter paper, and re-planted into pots. The transformed pepper plants were kept in the dark for 48 h before being transferred to growth conditions consisting of 26–28ºC,12 h light/12 h dark cycle, with their meristems sprayed with 50 mg/L kanamycin. We separated adventitious buds growing from kanamycin-resistant meristems when they reached a length of 1 cm and transplanted them to new pots containing peat soil to establish $T_0$ plants. These plants were further subjected to kanamycin, allowed to self-pollinate and set $T_1$ seeds. Seeds collected from $T_0$ plants were further screened by germinating them on MS medium supplemented with appropriate concentrations of 50 mg/L kanamycin. Resistant seedlings were confirmed by PCR-based genotyping using primers specific for the transgene (Table S1). We repeated the same procedure on seeds from $T_2$ and $T_3$ lines. All subsequent plant work was performed on homozygous $T_3$ lines.

Genetic transformation of *N. benthamiana* followed the method of Regner *et al.* (F *et al.*, 1992) and Bardonn *et al* (N *et al.*, 1994). Leaf discs of *N. benthamiana* were transformed with *Agrobacterium* cultures bearing various vectors (*35Spro:CaWRKY40-GFP* or *35Spro:CaIPT5-GFP*). Independent $T_0$ transgenic *N. benthamiana* plants were selected on the 10% PPT (glufosinate) and later confirmed by PCR with specific primers (Table S1). $T_0$ plants were allowed to self-pollinate and set seeds. Primary transformants were selected by germination on MS medium supplemented with PPT. Resistance status of seedlings was confirmed by PCR using specific primers (Table S1). We repeated the same procedure with seeds from the $T_2$ and $T_3$ lines. All subsequent plant work was performed on homozygous $T_3$ lines.

**Methods S3. Virus-Induced Gene Silencing (VIGS)**

VIGS assay was performed as previously described by Khan *et al.* (2018). We introduced the plasmids pTRV1, pTRV2:*00*, pTRV2:*CaPDS*, pTRV2:*CaWRKY40*, and pTRV2:*CaIPT5* into *Agrobacterium* strain GV3101. *Agrobacterium* cultures carrying each binary vector were then grown in 10 mL liquid LB medium with antibiotic selection (100 mg/mL kanamycin, 100 mg/mL rifampicin) at 28ºC overnight before collection by centrifugation and resuspension into *Agrobacterium* infiltration buffer (10 mM MES, 10 mM $MgCl_2$, 200 µM acetosyringone, pH 5.4) to an adjusted $OD_{600}$ = 0.8. Resuspended *Agrobacterium* cells carrying the pTRV1 vector were mixed in a 1:1 ratio with resuspended *Agrobacterium* cells bearing the pTRV2:*CaWRKY40*, pTRV2:*CaIPT5*, pTRV2:*00* or pTRV2:*CaPDS* binary vectors, incubated for 3 h on a shaker set at a low speed, and then co-infiltrated into the cotyledons of pepper seedlings.

Following infiltration, seedlings were kept at 16ºC in the dark for 56 h, and then returned to the growth chamber at 28ºC, 60–70 μmol photons $m^{-2}$ $s^{-1}$, a 16-h light/8-h dark photoperiod, and a relative humidity of 60%. Plants infiltrated with pTRV2:*CaPDS* provided an easily scorable visual confirmation of effective gene silencing, as evidenced by the appearance of bleached sectors due to silencing of *PHYTOENE DESATURASE* (*PDS*).

**Methods S4. Exogenous Applications of SA, JA, tZ, KT, or 6-BA**

Pepper, tomato and tobacco seedlings were grown in peat soil and kept in the growth chamber (27–29ºC, 60–70 μmol photons $m^{-2}$ $s^{-1}$, relative humidity of 60%, and 16-h light/8-h dark photoperiod) until the 6- to 8-leaf stage. Plant roots were then mechanically damaged were transferred to plastic cups filled with liquid MS medium containing 0.002 mM tZ, 0.002 mM KT, 0.002 mM 6-BA or $ddH_2O$, and inoculated with 1 mL of *R. solanacearum* cell suspension ($10^8$ cfu/mL). Pepper plants were then placed in an incubator with the temperature set to 30ºC, 34ºC, or 37ºC, 90% relative humidity, 16-h light/ 8-h dark photoperiod 15 days.

**Methods S5. RNA-seq Analysis**

Total RNA was extracted from the roots of pepper plants at the ten-leaf stage (inbred lines HN42 or TT5203) challenged with one of four treatments (RSHT, RSRT, RTHH, or HTHH) harvested at different time points. We also collected roots from *CaWRKY40*-silenced and wild-type pepper plants challenged with RSHT. Total RNA was extracted with the MagMAX™-96 Total RNA Isolation Kit (Ambion, AM1830) according to the manufacturer's instructions. mRNA sequencing libraries were constructed with barcodes using the TruSeq RNA Sample Preparation Kit (Illumina) with three biological replicates per sample, and sequenced on an Illumina HiSeq2500 platform by Novogene (Beijing, China) as 125 bp/150 bp paired-end reads. The RNA-seq data sets used in this study have been deposited at the CNSA repository(CNP0001156; CNP0001157).

We downloaded the reference genome and gene model annotation files from the pepper genome database (http://passport.pepper.snu.ac.kr/?t=PGENOME). We generated genome indices using Bowtie v2.2.3 and aligned quality-controlled and trimmed paired-end reads to the reference genome using TopHat v2.0.12. We obtained read counts per gene with HTSeq v0.6.1, which we then converted to fragments per kilobase of transcript per million mapped reads (FPKM) by normalizing read counts to gene length. We determined differential expression between two conditions/groups (three biological replicates per condition) using the DESeq R package (1.18.0). The resulting *P* values were adjusted using the Benjamini and Hochberg approach for controlling the false discovery rate. Genes with an adjusted *P* value <0.05 found by DESeq were deemed differentially expressed.

**Methods S6. EMSA Analysis**

We tested the *in vitro* binding of CaWRKY40 against the *cis*-elements W-box, WT, or ML or their mutant versions with synthesized DNA probes by EMSA. The positive-sense strand of each probe was labeled with the Cy5 fluorescent dye but the antisense strand was not labeled. Double-stranded wild-type and mutant probes (with or without Cy5 fluorescent label) were generated by mixing the positive-sense and antisense DNA strands and subjecting them to gradient freezing. Wild-type or mutant probes were mixed with GST or CaWRKY40-GST fusion protein and incubated in 5 × binding buffer (1 M Tris-HCl pH 7.5, 5 M NaCl, 1 M KCl, 1 M $MgCl_2$, 0.5 M EDTA pH 8.0, 10 mg/mL BSA) on ice for 30 min in the dark. The mixture was separated by PAGE gel and then scanned on an Odyssey$^{@}$ CLX fluorescence imaging system (LI-COR, USA).

**Methods S7. MST Analysis**

To analyze the binding of CaWRKY40 to DNA fragments containing the W- , WT-, or WL-box or their mutant versions in the context of the *CaIPT5* promoter *in vitro*, we performed an MST assay as previously described (Qiu *et al.*, 2018). The CaWRKY40-GFP fusion protein was isolated and purified from total protein extracts from pepper leaves transiently overexpressing *CaWRKY40-GFP* and served as fluorescent label. DNA fragments containing a W- , WT- , or WL-box or their mutant versions were used as ligand, diluted from 1.0E$^{-3}$ to 1.0E$^{-10}$ μM and then mixed with CaWRKY40-GFP fusion protein in interaction buffer (100 mM NaCl, 1 mM EDTA, 20 mM sodium phosphate, pH 8.0). The mixture was loaded into Monolith NT.115 Capillaries (Cat. #MO-K002, NanoTemper Technologies, Germany) using 50% IR laser power and an LED excitation source, where λ = 470 nm at ambient temperature. Dissociation constant ($K_d$) values were analyzed with the NanoTemper Analysis 1.2.20 software.

**Methods S8. GUS Activity Assay**

To assay the binding of CaWRKY40 to W-, WL-, and WT-box upon RTHH, RSRT, HTHH, or RSHT treatment and their possible effect, we constructed promoter reporter constructs, whereby the expression of the *GUS* gene was driven by synthetic promoters containing three copies of the W-, WL-, or WT-box or their corresponding mutant versions in the binary vector pMDC163. The pMDC163 vector and variants were introduced into *Agrobacterium* strain GV3101, and then infiltrated into the leaves of control or *CaWRKY40*-silenced pepper plants. After 24 h (post infiltration of hpi), all plants were challenged with RTHH, RSRT, HTHH, or RSHT treatment. Leaves were harvested and processed for GUS activity assessment 24 h later. We extracted total proteins in GUS protein extraction buffer (0.1 M Phosphate Buffered Saline (PBS) pH 7.0, 10% SDS, 0.5 M EDTA pH 8.0, 0.2% Triton X-100, 0.2% β-mercaptoethanol) and measured GUS activity using 4-methylumbelliferyl β-D-glucuronic acid (4-MUG) as a substrate according to Jefferson (JEFFERSON, 1987). We determined total protein concentration by the Bradford method (Bradford, 1976). To measure the concentration of the GUS product 4-methylumbelliferyl (4-MU) resulting from the conversion of 4-MUG, we added 5 μL total protein extract to 195 μL GUS assay buffer (0.44 mg/mL 4-MUG in GUS extraction buffer) and incubated the reactions at 37°C and measured 4-MU fluorescence (excitation: 365 nm; emission: 455 nm) after 10, 20, 30, and 60 min in a microplate reader (Synergy H1, biotek). GUS activity was expressed in units of pmol 4-MU produced per milligram of protein per minute.

**Methods S9. prokaryotic Expression and Protein Purification**

For expression of CaWRKY40 in *E. coli*, we introduced the pDEST15 vector containing a CaWRKY40-GST expression cassette into *E. coli* BL21 (DE3) strain. We induced the production of the CaWRKY40-GST recombinant protein by adding 100 μM isopropyl-β-d-thiogalactoside (IPTG) and allowing bacterial growth for 14 h at 20ºC. Bacteria were collected by centrifugation 7000 rpm, 5 min at 4ºC. Production of the target protein was confirmed by SDS-PAGE electrophoresis and Coomassie Brilliant Blue staining of the gel. To isolate and purify the fusion protein, the bacterial cells were resuspended in buffer A (140 mM NaCl, 2.7 mM KCl, 10 mM $Na_2HPO_4$, 1.8 mM $KH_2PO_4$, pH 7.4) by sonication with an Ultrasonic Cell Disruption System (JY96-IIN Xinzhi, China). Cell debris were pelleted by centrifugation at 4ºC, and the supernatant was mixed with BeaverBeads™ GSH (BEAVER, China) and then incubated with shaking for 3 h at 4ºC. Recombinant CaWRKY40-GST immobilized on BeaverBeads™ GSH was washed four times using buffer A (140 mM NaCl, 2.7 mM KCl, 10 mM $Na_2HPO_4$, 1.8 mM $KH_2PO_4$, pH 7.4) and then eluted with elution buffer B (50 mM Tris-HCl, 10 mM reduced glutathione, pH 8.0).

**Methods S10. plant Total Protein Extraction and Immunoblot Analysis**

We collected leaves from transgenic pepper or *N. benthamiana* plants, froze them in liquid nitrogen, and quickly ground the tissue into a fine powder, which was then mixed with protein extraction buffer (25 mM Tris-HCl pH 7.4, 150 mM NaCl, 1 mM EDTA, 10% glycerol, 2% Triton X-100, 2 mM DTT, 2% polyvinyl pyrrolidone, plant protease inhibitor cocktail). Extraction of total proteins was performed in 1.5 mL centrifuge tubes, slowly rotated at 4ºC for 1 h and then centrifuged at 4ºC, 20,000 *g* for 10 min. The supernatant was transferred to new cold tubes and kept on ice.

For immunoblot analysis, equal amounts of total protein were boiled in SDS loading buffer at 95ºC for 10 min and separated by SDS-PAGE. Separated proteins were then electro-transferred to PVDF membranes (Millipore, Cat.: IPVH00010) at a constant current of 200 mA for 30 min. Membranes were incubated in blocking buffer (1× Tris Buffered Saline, 0.05% Tween-20 (TBST) buffer + 5% skim milk powder [w/v]) for 1 h before incubation at room temperature with anti-GFP rabbit antibodies (MBL, sc-

9996) diluted 1:5,000 in blocking buffer. After three washed with TBST buffer, the anti-GFP antibody was detected using a peroxidase-conjugated goat anti-rabbit IgG (MBL, sc-2357).

**Methods S11. phytohormone Measurements and Quantification**

We determine the concentrations of the endogenous phytohormones SA, JA, ABA, cytokinins, IAA, and GA in the roots of pepper or transgenic *N. benthamiana* plants challenged with RSRT or RSHT treatment. We collected roots at the indicated time points, froze them in liquid nitrogen, and ground them to a fine powder. We weighted exactly 50 mg of each sample, to which we then added 0.5 mL of an isopropanol:water:hydrochloric acid (2:1:0.002,v:v:v) solution, as well as 10 ng $^{2}H_{2}$-GA3 and 20 ng each of $^{2}H_{5}$-IAA, $^{2}H_{6}$-ABA, $^{2}$H-JA, and $^{2}H_{5}$-SA at the same time as standards. The mixture was mixed by shaking for 30 min at 4ºC, and then 1 mL dichloromethane was added and shaking was continued for another 30 min. The lower layer solution was collected after centrifugation at 13,000 g for 5 min at 4ºC, dried under nitrogen flow, dissolved by adding 0.1 mL methanol, and centrifuged again at 13,000 g for 5 min at 4ºC. The final supernatant was analyzed by liquid chromatography tandem mass spectrometry (LC-MS/MS).

The chromatographic conditions were an Agilent SB-C18 column (4.6 mm × 50 mm, 1.8 μm), with mobile phase consisting of acetonitrile/0.1% acetic acid water and the following gradient elution: 0–0.5 min, 10% acetonitrile; 0.5–5 min, 10%–95% acetonitrile; 5–5.1 min, 95%–10% acetonitrile; 5.1–8.0 min, 10% acetonitrile. The flow rate was 0.8 mL/min, the column temperature was 30 ℃, and the injection volume was 10 μL. The mass spectrometry conditions were: electrospray ion source (ESI), negative ion scanning, multi-reaction monitoring, (MRM), air curtain gas pressure 30 psi, ionization voltage 4500 V, ion source temperature 600 ℃, spray gas pressure 40 psi, auxiliary gas pressure 50 psi. Data processing was performed with Analyst software.

### EXPERIMENTAL MODEL AND SUBJECT DETAILS

### KEY RESOURCES TABLE

| REAGENT or RESOURCE | SOURCE | IDENTIFIER |
|---|---|---|
| **Antibodies** | | |
| Anti-GFP (Green Fluorescent Protein) mAb | MBL | Cat. #D153-3 |
| Anti-GST-tag pAb | MBL | Cat. #PM013 |
| Phanta Max Super-Fidelity DNA Polymerase | Vazyme Biotech | Cat. #P505-d1 |
| Gateway™ BP Clonase™ II Enzyme mix | ThermoFisher | Cat. #11789020 |
| Gateway™ LR Clonase™ II Enzyme mix | ThermoFisher | Cat. #11791100 |
| Anti-Histone H3 (di methyl K9) antibody [mAbcam 1220] - ChIP Grade | abcam | Cat. #ab1220 |
| Anti-Histone H3 (tri methyl K4) antibody - ChIP Grade | abcam | Cat. #ab8580 |
| **Bacterial Strains** | | |
| *Agrobacterium tumefaciens* GV3101 | N/A | N/A |
| *Escherichia coli* Rosetta 2(DE3) | Weidi Biotechnology | Cat. #EC1014 |
| *Escherichia coli* DH5α | Weidi Biotechnology | Cat. #DL1001 |
| *Ralstonia solanacearum* GMI1000 | N/A | N/A |
| **Chemicals, Peptides, and Recombinant Proteins** | | |
| GFP-Trap®_A (for immunoprecipitation) | Chromotech | Cat. #gta-20 |
| BeaverBeads™ GSH | Beaver Biosciences | Cat. #70601-5 |
| TRIzol Reagent | ThermoFisher | Cat. #15596026 |
| Protease inhibitor cocktail | Sigma-Aldrich | Cat. #P9599-5ML |
| Salicylic acid, SA | Sigma | Cat. #247588 |
| Jasmonic acid, JA | Sigma | Cat. #77026-92-7 |
| Abscisic acid, ABA | Sigma | Cat. #A 1049 |
| Zeatin, ZT | Sigma | Cat. #32771-64-5 |
| *trans*-Zeatin, tZ | Sigma | Cat. #1637-39-4 |
| Kinetin, KT | Sigma | Cat. #525-79-1 |
| 6-Benzylaminopurine, 6-BA | Sigma | Cat. #1214-39-7 |
| Indoleacetic acid , IAA | Sigma | Cat. #87-51-4 |
| GST-CaWRKY40 | This paper | N/A |
| Critical Commercial Assays | | |
| HiScript Q RT SuperMix for qPCR | Vazyme Biotech | Cat. #R122-01 |
| AceQ qPCR SYBR Green Master Mix | Vazyme Biotech | Cat. #Q111-02 |
| **Deposited Data** | | |
| Raw and analyzed data | This paper | CNP0001155; CNP0001156; CNP0001157 |
| *Capsicum annuum* genome | The pepper Information Resource | http://passport.pepper.snu.ac.kr/?t=PGENOME |
| **Experimental Models: Organisms/Strains** | | |

| | | |
|---|---|---|
| Pepper: 35Spro:CaWRKY40-GFP | This paper | N/A |
| *Nicotiana benthamiana*: 35S:CaWRKY40-GFP | This paper | N/A |
| *Nicotiana benthamiana*: 35S:CaIPT5-GFP | This paper | N/A |
| **Recombinant DNA** | | |
| pEG103-35S-CaWRKY40-GFP | This paper | N/A |
| pEG103-35S-CaIPT5-GFP | This paper | N/A |
| pDEST15-T7-GST-CaWRKY40 | This paper | N/A |
| pTRV2-CaWRKY40 | This paper | N/A |
| pTRV2-CaIPT5 | This paper | N/A |
| pTRV1 | This paper | N/A |
| **Sequence-Based Reagents** | | |
| For primers used in this study, see Table S1 | This paper | N/A |
| **Software and Algorithms** | | |
| DPS 7.05 | N/A | N/A |
| Sigmaplot 12.5 | Systat | N/A |
| NanoTemper Analysis 1.2.20 | N/A | N/A |

### Tables

### Table S1. Primers used in this study.

| | Gene | Zunla genome ID or NCBI Address | F_primer | R_primer |
|---|---|---|---|---|
| Primers used for *CaWRKY40* and *CaIPT5* study | ***CaWRKY40***[1] | Capana06g001110 | CGGAATTCATGATGGAATTTACCAGTTTGGTTGA | CGGGATCCTTACCATCTGCCCGTCTGA |
| | ***CaWRKY40*-GFP**[2] | Capana06g001110 | CGGAATTCATGATGGAATTTACCAGTTTGGTTGA | CGGGATCCCCATCTGCCCGTCTGA |
| | ***CaWRKY40*-VIGS**[3] | Capana06g001110 | AAAAAGCAGGCTTCAGAAGAGCTTTGTTGATTCT3 | GAGAAAGCTGGGTCACCATTTTGTGTGCTCATAT |
| | ***CaIPT5-GFP***[2] | Capana01g003019 | GGGGACAAGTTTGTACAAAAAAGCAGGCTTCGGATGATCAATATTGCACCATT | GGGGACCACTTTGTACAAGAAAGCTGGGTCGTGCGTTGCAGTAGCAA |
| | ***CaIPT5*-VIGS**[4] | Capana01g003019 | GGGGACAAGTTTGTACAAAAAAGCAGGCTTCGGTGGATTGATGTGAAGTTGGA | GGGGACCACTTTGTACAAGAAAGCTGGGTCTTGAGGAAGAAGATGGAGTAG |
| Primers used for pepper qPCR analysis | ***CaWRKY40*-qPCR** | Capana06g001110 | AACTTGGATGTTGTGCCTGGA | CTGTAACCTTGGCTTTTATGTGC |
| | ***CaIPT5*-qPCR** | Capana01g003019 | CGTCGGCTTGTATCTGAA | GTAAGAAGGAAGTTGAGGAAG |
| | ***CaICS1*-qPCR** | Capana06g001004 | TGGTCAGTGTTGCTGGTGTT | GCATCAAATCGGATTGCCCC |
| | ***CaSTH2*-qPCR** | Capana03g004445 | ACACACCCTAAAATCTCACTCCA | GCCTTAAACAAGAAGAAGAAAGAGCT |
| | ***CaDEF1*-qPCR** | Capana07g000107 | TCTTTGCTCGTAATGATTTGTGACA | GGTGTTCTTGGCTTATAGTGGC |
| | ***CaMgst3*-qPCR** | Capana02g002296 | TCCAGCTTTTCGCACTCTCT | CGAGATCTCGCCACCCAATT |
| | ***CaPRP1*-qPCR** | Capana09g001760 | ACCAAACTCGGATATTCCAACT | GAGGAATCCTCGGAACCAAGT |
| | ***CaACTIN*-qPCR** | GQ339766 | AGGGATGGGTCAAAAGGATGC | GAGACAACACCGCCTGAATAGC |
| | **GFP-qPCR**[5] | | AGGAGCGCACCATCTTCTTC | AAGTCGATGCCCTTCAGCTC |

| | Gene | Zunla genome ID or NCBI Address | F_primer | R_primer |
|---|---|---|---|---|
| Primers used for *Nb* qPCR analysis | **NbWRKY40-qPCR** | Niben101Scf00715g09001 | ACTGCTTCAGTCCCTTGCTC | CCTTCTGTTGTCCTGTGGCT |
| | ***NbIPT5*-qPCR** | Niben101Scf04783Ctg038 | GCATCAAGCTTGGCATAGGC | TGCAGGAGGTTTGGTGGTTT |
| | ***NbICS*-qPCR** | Niben101Scf05166g06006. | ACCCTTTGCCGGAAGAAGAG | GCCTCTTGTCAGAGCCTGTT |
| | ***NbSTH2*-qPCR** | Niben101Scf10735g00016 | TCACAACATCAGCTTCCCCA | CCCCCATCACCCTCAACAAT |
| | ***NbDEF1*-qPCR** | Niben101Scf06275g03010 | GCCTTACCAAACCACCATGC | GCTGCAGCCAAAGTTTTTGC |
| | ***NbMgst3*-qPCR** | Niben101Scf07253g02012 | CAAAACCCAAATGGCCGGAG | CTTTTTGCGAGCTTTGCCGA |
| | ***NbPRP1*-qPCR** | Niben101Scf10316g03002 | GCTTGGTGTTTCACTCAGCC | GCAGAGGGCTCTTGTTTTGT |
| | ***NtEF-1a*-qPCR** | D63396 | TGCTGCTGTAACAAGATGGATGC | GAGATGGGGACAAAGGGGATT |
| Primers used for ChIP-PCR analysis | ***pCaIPT5-Wl*** | Capana01g003019 | AAGTATATATTCATGGACTC | GTTACAAAAGACCACGTAAT |
| | ***pCaIPT5-CK*** | Capana01g003019 | CTGGTTTAAAGCCAGTTCAAGT | TGTTAACAATGTGGAGTGGTGT |
| | ***pCaICS1-Wt*** | Capana06g001004 | CTAACGCCCCTTTGCCTGTA | ACTCTCTCGTCGTCTTGTGC |
| | ***pCaICS1-CK*** | Capana06g001004 | TGTTTGCAACCACGGTGTTT | TGGGTTCCAAAGTACACGGA |
| | ***pCaSTH2-Wb1*** | Capana03g004445 | ATCTGATCCTTGATTATTGC | AATTTATTATGTGCATGACA |
| | ***pCaSTH2-Wb2*** | Capana03g004445 | AAATTTGTTTTTCTAAGAGA | GGATAAGATGATTGTAATAT |
| | ***pCaSTH2-CK*** | Capana03g004445 | TGCAAACTCATGAGACTGTAACT | AGTTGATGTAGACTGTGGGAAC |
| | ***pCaDEF1-Wb*** | Capana07g000107 | GCTATAAAACCATCATATTG | TAGTGTACGTGTAGGACCAT |
| | ***pCaDEF1-Wl*** | Capana07g000107 | CGTTGATGAGCAAAAGTAGCGA | AGACTTTTGAATGATGTTTGTGGT |
| | ***pCaDEF1-CK*** | Capana07g000107 | CCTCCGGAATCATGTCACCC | AGGTTAAACATTTTGGACCATTTGGA |
| | ***pCaMgst3-Wb1*** | Capana02g002296 | TCCACTCTTCCACTTGACTT | CAATCTTCGGTTGAAGCAAT |
| | ***pCaMgst3-Wb2*** | Capana02g002296 | GTTGCGAGTGACATGTTTAT | GTTCTTGTTAGTGCAGTACT |
| | ***pCaMgst3-Wl*** | Capana02g002296 | AGATCTCGAACCAGTCCCCT | GGCTTCTTGATCTTTCTGAGAGA |
| | ***pCaMgst3-CK*** | Capana02g002296 | GGCAAAAACAAGAAGGGTTTTCC | TGGTTCTGCTATCAAGAACGA |
| | ***pCaPRP1-Wt*** | Capana09g001760 | TATGACCTCCCTCCAAGTCAG | TTGTAACCCCCATGCGTGAG |
| | ***pCaPRP1-Wl*** | Capana09g001760 | TCGGAGGTCATATTTGTTCAAATTT | AGTTGGAATATCCGAGTTTGGT |
| | ***pCaPRP1-Wb*** | Capana09g001760 | GATATTAATAAGTAAAAATG | TTTATTTCAGTATGTGTTGG |
| | ***pCaPRP1-CK*** | Capana09g001760 | AGTCGAAAATTATTTTCCGACAAGC | GCCAATGTAGTCGGAATGTGG |
| | ***CaIPT5-Tss*** | Capana01g003019 | CCAAAAATCAGGCTCGCTATATCT | ACGGACCTGTTAAACAAGAAGGA |
| | ***CaICS1-Tss*** | Capana06g001004 | TGTTTGCAACCACGGTGTTT | TGGGTTCCAAAGTACACGGA |
| | ***CaSTH2-Tss*** | Capana03g004445 | ACACACCCTAAAATCTCACTCCA | GCCTTAAACAAGAAGAAGAAAGAGCT |
| | ***CaDEF1-Tss*** | Capana07g000107 | TCTTTGCTCGTAATGATTTGTGACA | GGTGTTCTTGGCTTATAGTGGC |

| | | | | |
|---|---|---|---|---|
| | ***CaMgst3-Tss*** | Capana02g002296 | TCCAGCTTTTCGCACTCTCT | CGAGATCTCGCCACCCAATT |
| | ***CaPRP1-Tss*** | Capana09g001760 | TGACTTGGTTCCGAGGATTCC | CTCACAACTTTGGTTCTCTGTTT |

[1]Primers used for *CaWRKY40* full-length cloning

[2]Primers used for *35Spro:CaWRKY40-GFP* and *35Spro:CaIPT5-GFP* construct

[3]Primers used for pTRV:*CaWRKY40* construct

[4]Primers used for pTRV:*CaIPT5* construct

[5]Primers used for transgenic *N. benthamiana* and pepper plants test

**Table S2. Disease index for pepper plants infected with *R. solanacearum***

| **Score** | **Condition level** |
|---|---|
| 0 | Pepper plants are normal and asymptomatic. |
| 1 | Plants show slight withering, <25% leaves are withered, the apical region of the plant is normal. |
| 2 | Leaves in addition to the top leaves are affected, 25–50% of leaves are withered, the apical region of the plant is normal. |
| 3 | 50–75% of leaves are withered, while the top of the plant is normal. |
| 4 | >75% of leaves are withered, or plant is dead. |

**Table S3. Venn diagram results for Figure 4C**

| Names | total | elements |
|---|---|---|
| CaWRKY40-induced *PRs* after RSHT vs HN42-induced *PRs* after RSHT vs *PRs* in CaWRKY40 targets | 2 | Capana09g001760<br>Capana02g002296 |
| CaWRKY40-induced *PRs* after RSHT vs HN42-induced *PRs* after RSHT | 16 | Capana10g002344 Capana03g004566<br>Capana00g003098 Capana09g001761<br>Capana07g002003 Capana00g000744<br>Capana09g001860 Capana00g002164<br>Capana03g004562 Capana00g003222<br>Capana00g003097 Capana12g000240<br>Capana02g000952 Capana08g002190<br>Capana03g004199<br>Capana07g002012 |
| CaWRKY40-induced *PRs* after RSHT vs *PRs* in CaWRKY40 targets | 1 | Capana03g004445 |
| CaWRKY40-induced *PRs* after RSHT | 15 | Capana05g001164 Capana00g004929<br>Capana03g004443 Capana00g004920<br>Capana02g002478 Capana08g002823<br>Capana09g001741 Capana08g001910<br>Capana04g000545 Capana02g002479<br>Capana11g001524 Capana09g001763<br>Capana03g004442 Capana03g004448<br>Capana03g004407 |
| HN42-induced *PRs* after RSHT | 14 | Capana07g000035 Capana03g001052<br>Capana04g002897 Capana02g001373<br>Capana11g001537 Capana03g001050<br>Capana00g003106 Capana09g001740<br>Capana08g001518 Capana12g000354<br>Capana11g001535 Capana11g000674<br>Capana06g000760<br>Capana03g000768 |
| *PRs* in CaWRKY40 targets | 9 | Capana07g000107 Capana03g004402<br>Capana00g003023 Capana12g002446<br>Capana00g003438 Capana00g000196<br>Capana07g000894 Capana11g001711<br>Capana08g001755 |

**Figures and Legends**

**Figure S1. Phenotypic Response of Plants from Different Inbred Pepper Lines to RSRT, HTHH, or RSHT Treatment**

(a) Phenotypic response of different pepper inbred lines to RSRT, HTHH, or RSHT treatment at 3 days post treatment (dpt).

(b) Growth of *R. solanacearum*, as determined by colony forming units (cfu) detected in leaves from different *R. solanacearum*-inoculated pepper inbred lines challenged by RTHH or HTHH treatment at 24, 48, and 72 h post inoculation (hpi), which was carried out by root irrigation.

(c) Phenotypic response of different pepper inbred lines to RSHT treatment at 7 dpt.

**Figure S2. Phenotypic Response of *N. benthamiana* and Tomato (*Solanum lycopersicum*) Plants to RSRT, HTHH, or RSHT Treatment**

(a) Phenotypic response of *N. benthamiana* and tomato plants to RSRT, HTHH, or RSHT treatment at 3 dpt.

(b) Survival rate of *N. benthamiana* and tomato plants to RSRT, HTHH, or RSHT treatment, scored from 0 to 18 dpt. A total of 50 plants were dynamically scored for each species.

**Figure S3. Heatmap Representation of the Transcript Levels of Putative R Genes upon RTHH, RSRT, and RSHT Treatment in the TT5203 and HN42 Pepper Inbred Lines**

Transcript levels are displayed using a color scale in which blue, black, and yellow indicate low, intermediate, and high levels, respectively. Group number is indicated at left.

**Figure S4. Transcript Levels of Members of the Pepper *WRKY* Gene Family upon RTHH, RSRT, HTHH, or RSHT Treatment**

(a) Transcript levels of pepper *WRKY* family members in the roots of TT5203 and HN42 plants challenged with RTHH, RSRT, HTHH, or RSHT treatment, visualized as a heatmap. Blue, black, and yellow indicate low, intermediate, and high levels, respectively.

(b) Number of unchanged, up-regulated, and down-regulated *WRKY* genes in the roots of TT5203 and HN42 plants challenged with RSRT, HTHH, or RSHT treatment. Total number of differentially expressed *WRKY* genes is given in parentheses.

(c) Venn diagram of the number of *WRKY* genes up-regulated in roots of HN42 and TT5203 pepper plants challenged with RSHT treatment. The three *WRKY* genes specifically up-regulated in the HN42 inbred line are listed at right. *CaWRKY40* exhibited the highest transcript levels and is labeled in red.

(d) Expression profile of *CaWRKY40* in HN42 pepper plant roots upon RT, RSRT, HT, or RSHT treatment at 1, 3, 6, 12, 24, and 48 hours post treatment (hpt). The *CaACTIN1* was used as an internal control. Data represent the mean ± SD of three biologically replicates. Different capital letters above the bars indicate significant difference ($P <$ 0.01), determined using Fisher's protected LSD test.

**Figure S5. The Contents of the Phytohormones SA, ZT, tZ, ABA, and JA-Ile in Roots of *N. benthamiana* Plants Overexpressing *CaWRKY40* and Their Control Plants after Challenge with RSRT or RSHT Treatment at 48 hpt**
Data represent the mean ± SD of three biological replicates. Different capital letters above the bars indicate significant difference ($P$ < 0.01), determined using Fisher's protected LSD test.

**Figure S6. Binding of CaWRKY40 to the *STH2, DEF1, Mgst3*, and *PRP1* Promoters by ChIP-PCR Analysis**
(a) Integrative Genomics Viewer (IGV) images of ChIP-seq data and the location of W-box, WT, and WL motifs within the promoters of *STH2*, *DEF1*, *Mgst3*, and *PRP1* genes.
(b) Accumulation of CaWRKY40 in leaves of pepper plants overexpressing CaWRKY40-GFP and challenged with RTHH, RSRT, HTHH, or RSHT treatment, by immunoblot analysis. α-GFP, anti-GFP antibodies; CBB, Coomassie Brilliant Blue.
(c) Binding of *Ca*WRKY40-GFP to W-box, WT, and WL motifs within the promoters of *STH2*, *DEF1*, *Mgst3*, and *PRP1*, by ChIP-PCR analysis. Roots from *CaWRKY40-GFP* transgenic pepper plants treated with RTHH, RSRT, HTHH, or RSHT were harvested for ChIP-PCR analysis at 48 hpt.

**Figure S7. IPT5-mediated Immunity Participates in Pepper Responses to RSHT but not to RSRT**
(a) Relative transcript levels of *CaWRKY40*, *STH2*, *DEF1*, *Mgst3*, *PRP1*, and *ICS1* in roots of pTRV:*IPT5* and control pepper plants challenged with RTHH, RSRT, HTHH, or RSHT treatment, as determined by RT-qPCR at 48 hpt. *CaACTIN* was used as an internal control.
(b) Relative transcript levels of *NbWRKY40*, *NbSTH2*, *NbDEF1*, *NbMgst3*, and *NbPRP1* in roots of *N. benthamiana* plants overexpressing *CaIPT5* and their control plants challenged with RTHH, RSRT, HTHH, or RSHT, as determined by RT-qPCR at 48 hpt. *NbACTIN* was used as an internal control.
In (a) and (b), data represent the mean ± SD of three biological replicates. Different capital letters above the bars indicate significant difference ($P$ < 0.01), determined using Fisher's protected LSD test.

**Figure S8. The Phytohormone Content for ABA, JA-Ile, SA, tZ, and ZT in Roots of *CaIPT5*-silenced HN42 and Control Pepper Plants Challenged with RTHH, RSRT, HTHH, or RSHT Treatment at 48 hpt**
Data represent the mean ± SD of three biological replicates. Different capital letters above the bars indicate significant difference ($P$ < 0.01), determined using Fisher's protected LSD test.

**Figure S9. The Effect of Exogenous Application of the Phytohormones SA, MeJA, and tZ on the Response of Pepper, Tomato and Tobacco Plants to RSRT and RSHT Treatments**

(a) Effect of exogenously applied SA, MeJA, or tZ on phenotypic responses of pepper, tomato, and tobacco plants to RSI under RTHH or HTHH (28–37ºC, high relative humidity).

(b) The effect of exogenous SA, MeJA, or tZ on diseases index of pepper, tomato, and tobacco plants from 0 to 15 dpt upon inoculation with R. solanacearum under RTHH or HTHH (28ºC, 31ºC, 34ºC, or 37ºC, high relative humidity). A total of 12 pepper, tomato, and tobacco plants each were dynamically scored.

**Bradford MM. 1976.** A Rapid and Sensitive Method for the Quantitation of Microgram Quantities of Protein Utilizing the Principle of Protein-Dye Binding. *Analytical Biochemistry* **72**(1-2): 248-254.

**F R, A dCM, ML dCM, H S, D M, V H, H W, H K. 1992.** Coat protein mediated resistance to Plum Pox Virus in Nicotiana clevelandii and N. benthamiana. *Plant cell reports* **11**(1): 30-33.

**JEFFERSON RA. 1987.** GUS fusion : β-glucuronidase as a sensitive and versatile gene fusion marker in higher plants. *Embo Journal* **6**.

**Khan MI, Zhang Y, Liu Z, Hu J, Liu C, Yang S, Hussain A, Furqan Ashraf M, Noman A, Shen L, et al. 2018.** *CaWRKY40b* in pepper acts as a negative regulator in response to *Ralstonia solanacearum* by directly modulating defense genes including *CaWRKY40*. *International Journal of Molecular Sciences* **19**(5).

**N B, F H, MA S, L P. 1994.** Protection against virus infection in tobacco plants expressing the coat protein of grapevine fanleaf nepovirus. *Plant cell reports* **13**(6): 357-360.

**Qiu A, Lei Y, Yang S, Wu J, Li J, Bao B, Cai Y, Wang S, Lin J, Wang Y. 2018.** CaC3H14 encoding a tandem CCCH zinc finger protein is directly targeted by CaWRKY40 and positively regulates the response of pepper to inoculation by Ralstonia solanacearum. *Molecular plant pathology* **19**(10): 2221.

**F R, A dCM, ML dCM, H S, D M, V H, H W, H K. 1992.** Coat protein mediated resistance to Plum Pox Virus in Nicotiana clevelandii and N. benthamiana. *Plant cell reports* **11**(1): 30-33.

**Khan MI, Zhang Y, Liu Z, Hu J, Liu C, Yang S, Hussain A, Furqan Ashraf M, Noman A, Shen L, et al. 2018.** *CaWRKY40b* in pepper acts as a negative regulator in response to *Ralstonia solanacearum* by directly modulating defense genes including *CaWRKY40*. *International Journal of Molecular Sciences* **19**(5).

**N B, F H, MA S, L P. 1994.** Protection against virus infection in tobacco plants expressing the coat protein of grapevine fanleaf nepovirus. *Plant cell reports* **13**(6): 357-360.

**F, R., A, d.C.M., ML, d.C.M., H, S., D, M., V, H., H, W., and H, K**. (1992). Coat protein mediated resistance to Plum Pox Virus in Nicotiana clevelandii and N. benthamiana. Plant cell reports *11*, 30-33.

**Khan, M.I., Zhang, Y., Liu, Z., Hu, J., Liu, C., Yang, S., Hussain, A., Furqan Ashraf, M., Noman, A., Shen, L.,** ***et al.*** **(2018).** *CaWRKY40b* in pepper acts as a negative regulator in response to *Ralstonia solanacearum* by directly modulating defense genes including *CaWRKY40*. International Journal of Molecular Sciences *19*.

**N, B., F, H., MA, S., and L, P. (1994).** Protection against virus infection in tobacco plants expressing the coat protein of grapevine fanleaf nepovirus. Plant cell reports *13*, 357-360.
